# Capture-based enrichment of *Theileria parva* DNA enables full genome assembly of first buffalo-derived strain and reveals exceptional intra-specific genetic diversity

**DOI:** 10.1101/2020.04.11.037309

**Authors:** Nicholas C Palmateer, Kyle Tretina, Joshua Orvis, Olukemi O Ifeonu, Jonathan Crabtree, Elliott Drabék, Roger Pelle, Elias Awino, Hanzel T Gotia, James B Munro, Luke Tallon, W Ivan Morrison, Claudia A Daubenberger, Vish Nene, Donald P Knowles, Richard P Bishop, Joana C Silva

**Affiliations:** Institute for Genome Sciences, University of Maryland School of Medicine, Baltimore, MD 21201, USA; Biosciences eastern and central Africa-International Livestock Research Institute, Nairobi, Kenya; International Livestock Research Institute, Nairobi, Kenya; The Roslin Institute, Royal (Dick) School of Veterinary Studies, University of Edinburgh, Easter Bush Campus, Roslin, Midlothian EH25 9RG, UK; Swiss Tropical and Public Health Institute, 4002 Basel, Switzerland; University of Basel, Basel, Switzerland; Department of Veterinary Microbiology and Pathology, Washington State University, Pullman, WA 99163, USA; Department of Microbiology and Immunology, University of Maryland School of Medicine, Baltimore, MD 21201, USA

**Keywords:** *Theileria parva*, lawrencei, DNA enrichment, genome assembly, polymorphism

## Abstract

*Theileria parva* is an economically important, intracellular, tick-transmitted parasite of cattle. A live vaccine against the parasite is effective against challenge from cattle-transmissible *T. parva* but not against genotypes originating from the African Cape buffalo, a major wildlife reservoir, prompting the need to characterize genome-wide variation within and between cattle- and buffalo-associated *T. parva* populations. Here, we describe a capture-based target enrichment approach that enables, for the first time, *de novo* assembly of nearly complete *T. parva* genomes derived from infected host cell lines. This approach has exceptionally high specificity and sensitivity and is successful for both cattle- and buffalo-derived *T. parva* parasites. *De novo* genome assemblies generated for cattle genotypes differ from the reference by ∼54K single nucleotide polymorphisms (SNPs) throughout the 8.31 Mb genome, an average of 6.5 SNPs/kb. We report the first buffalo-derived *T. parva* genome, which is larger than the genome from the reference, cattle-derived, Muguga strain. The average non-synonymous nucleotide diversity (*π*_N_) per gene, between buffalo-derived *T. parva* and the Muguga strain, was 1.3%. This remarkably high level of genetic divergence is supported by an average *F*_ST_, genome-wide, of 0.44, reflecting a degree of genetic differentiation between cattle- and buffalo-derived *T. parva* parasites more commonly seen between, rather than within, species, with clear implications for vaccine development. The DNA capture approach used provides clear advantages over alternative *T. parva* DNA enrichment methods used previously and enables in-depth comparative genomics in this apicomplexan parasite.

## Introduction

In developing countries, infectious diseases of livestock can have a broad and profound negative effect on public health, including malnutrition, increased susceptibility to disease, female illiteracy and loss of productivity (Rist et al. 2015; Thumbi et al. 2015; Marsh et al. 2016), and curb the potential for economic improvement (Black et al. 2013). *Theileria parva* is a tick-transmitted, obligate intracellular apicomplexan parasite that causes East Coast fever (ECF), an acute fatal disease of cattle in eastern, central and southern Africa. During proliferation of *T. parva* in the mammalian host, a multi-nucleated schizont immortalizes infected host lymphocytes and divides in synchrony with them, ensuring that the infection is transmitted to each daughter cell, through poorly understood mechanisms (von Schubert et al. 2010; Tretina et al. 2015; Marsolier et al. 2019; Huber et al. 2020; Tretina et al. 2020a). Susceptible animals usually die within three to four weeks post-infection. This is a result of widespread lysis of infected and uninfected lymphocytes in the lymphoid tissues, secondarily inducing a severe macrophage response, characterized by high IL-17 expression, and pulmonary edema (Fry et al. 2016). ECF represents a severe economic constraint and is a major impediment to the development of the cattle industry in the impacted region, affecting millions of cattle and killing over a million animals each year (Norval et al. 1991; Okuthe and Buyu 2006; Bazarusanga et al. 2007).

An infection-and-treatment method (ITM) involving administration of a lethal dose of a cryopreserved stabilate of *T. parva* sporozoites from three parasite isolates, together with a long-acting formulation of oxytetracycline, has been in use for several decades. Incorporation of the three parasite isolates (known as the Muguga cocktail) is required to avoid vaccine evasion, a common problem of anti-parasitic vaccines (Thera et al. 2011; Neafsey et al. 2015; Bishop et al. 2020). This vaccination method can protect against ECF for at least 43 months, although, in a variable percentage of animals, heterologous parasites may induce transient clinical symptoms in vaccinated cattle (Burridge et al. 1972). However, ITM also has significant drawbacks, including a logistically intensive manufacturing process (reviewed in Di Giulio et al. 2009). Also, there have been recently verified concerns that ITM is not always as effective against challenge from buffalo-derived *T. parva* as it is against cattle-derived parasites (Sitt et al. 2015), and even cattle-derived *T. parva* from geographically diverse regions could sometimes break through immunity induced by the Muguga cocktail vaccine (Amzati et al. 2019). The African buffalo (*Syncerus caffer*) is an asymptomatic wildlife carrier of *T. parva* in the region and is the primary mammalian host (Conrad et al. 1987). Areas where buffalo and cattle co-graze enable transmission of the parasite between mammalian hosts by the tick vector, *Rhipicephalus appendiculatus* (Bishop et al. 2015).

Studies using a limited set of markers strongly suggest that the *T. parva* strains (or genotypes) circulating in the affected cattle population represent only a subset of a much more heterogeneous *T. parva* meta-population residing in buffalo (Toye et al. 1995; Bishop et al. 2002; Oura et al. 2011; Pelle et al. 2011), due primarily to lack of tick transmissibility of buffalo-derived infections, associated with very low piroplasm counts. *T. parva* isolates obtained from buffalo were at one time classified into a separate subspecies, *T. parva lawrencei*, based on clinical presentation, despite the lack of genetic evidence to support this claim (Norval et al. 1991). Preliminary data suggests that genome-wide differences between the reference Muguga strain and buffalo-derived isolates are substantially larger than among cattle-transmissible genotypes (Hayashida et al. 2013), although there is currently no genome assembly for *T. parva* from buffalo. The design of a vaccine that is effective against most cattle- and buffalo-derived *T. parva* requires the comprehensive characterization of genetic differences within and between those two *T. parva* parasite populations, particularly in regions of the genome that encode antigenic proteins. Comprehensive knowledge of genetic variation in the species is also needed to monitor the impact of live vaccination on the composition of parasite field populations.

The biology of *T. parva* has so far proved a powerful obstacle to the acquisition of DNA in sufficient quantity and quality for whole genome sequencing. DNA extracted from cattle blood early in the infection cycle is heavily contaminated with host DNA. In late stages of infection, the tick-infective piroplasm stage infects erythrocytes, and requires collection of large volumes of blood from clinically ill *T. parva-*infected animals to obtain purified piroplasm DNA in sufficient quantity for genome sequencing (Gardner et al. 2005), an approach not sustainable or ethically feasible for a large number of strains. In addition, and despite their higher virulence to cattle, *T. parva* of buffalo origin induce lower levels of schizont parasitosis, and produce no or very few piroplasms in cattle (Norval et al. 1992), precluding their use as a source of parasite DNA. *T. parva* DNA can also be obtained from schizonts purified following lysis of infected lymphoblasts (Sugimoto et al. 1988; Goddeeris et al. 1991) but low yield, host DNA contamination, and the heterogeneity in lysing properties of infected cells make this approach unsuitable for high throughput applications.

Finally, the estimation of genome-wide population genetic diversity relies on the identification of sequence variants from the alignment of whole genome sequence data to a reference genome (Hedrick 2011). The same approach has been used to identify pathogen-encoded antigens, which are potential vaccine candidates, since they are often among the most variable protein-coding genes in a genome (Amambua-Ngwa et al. 2012). However, in highly polymorphic species such as *T. parva*, this approach is unreliable because sequence reads fail to map between strains, particularly in the genomic regions that encode the most variable antigens (Hayashida et al. 2013; Norling et al. 2015).

Here, we applied a target DNA sequence capture approach to selectively enrich parasite DNA in samples obtained from *T. parva*-infected bovine lymphocyte cultures, consisting mostly of bovine DNA (Gotia et al. 2016). Even though conceptually similar to pathogen DNA enrichment approaches used before for other organisms (Bright et al. 2012; Feehery et al. 2013; Christiansen et al. 2014; Huang et al. 2018), design choices in the current study resulted in extremely high capture sensitivity and specificity. Furthermore, to gain access to variable genomic regions that cannot be analyzed through read mapping approaches (Norling et al. 2015), we assembled the captured sequence read data and analyzed the resulting *de novo* genome assemblies for completeness. Starting from cell cultures in which the parasite DNA was less than 4% (Gotia et al. 2016), we have generated *de novo* genome assemblies for each isolate consisting of 109 - 126 scaffolds, that encompass >95% of the reference genome of *T. parva*. This approach was successful even when applied to a highly divergent *T. parva* isolate from buffalo, for which we present the first publicly available genome assembly. The ability to characterize genome-wide polymorphism based on whole genome assemblies, which provide higher resolution relative to read mapping approaches, particularly in highly variable regions of the genome, represents a powerful approach for the characterization of genetic variation in intracellular parasites such as *Theileria*, and in particular for the study of highly polymorphic antigens and other variable genes and regions of the genome.

## Results

### Design of the whole-genome sequence capture approach: Length and genome coverage of the probe set, and genomic library fragment size

We have customized a DNA sequence capture approach to obtain *T. parva* genomic DNA from *T. parva*-infected bovine lymphocyte cell lines. The premise is similar to that of exome capture (Hodges et al. 2007; Gnirke et al. 2009) in that the target DNA is only a small subset of the total DNA mix, but here the DNA fraction intended for capture is the 8.31 Mb-long *T. parva* nuclear genome and the 39 kb-long apicoplast genome, while the non-target DNA is the >300 times larger animal host genome. The capture probe set targets almost completely span the nuclear and apicoplast genomes of the reference *T. parva* Muguga (Gardner et al. 2005), and are based on the SeqCap EZ platform (Roche/Nimblegen). The probe design was conducted by Roche/Nimblegen using proprietary software. As part of the probe design, probe length was minimized, to increase the success of cross-strain DNA capture of loci with highly diverged segments by taking advantage of relatively small, conserved DNA segments that are intermixed with more rapidly evolving regions; the resulting probes average 76 bp in length. Probes that mapped to low complexity sequences or >5 genomic regions were eliminated, as were those with strong sequence similarity to the bovine genome. The final probe set consists of a series of overlapping probes, which cover 7,932,549 bp (95.5%) of the combined length of the nuclear genome (8,308,027 bp) and the apicoplast genome (39,579 bp). The fraction of the two genomes not covered by probes is spread among 3,843 independent genomic regions that average 93 bp in length (**Figure 1, Supplemental Table S1**). In total, 53 genes have no probe coverage, while 4,111 genes have at least some coverage by the probe set, with >50% of all genes being completely covered by probes (**Supplemental Table S1**). The 53 genes without probe coverage are all members of multigene families, including *Theileria parva* repeat (*Tpr*) and Subtelomere-encoded Variable Secreted Protein (SVSP). To maximize the probability of capturing both genes that are highly variable across strains and genomic regions without probe coverage, we sheared the genomic DNA sample to a fragment size significantly larger than the length of the probes, such that captured DNA fragments contain both conserved segments and highly variable regions that flank them.

**Figure 1.**
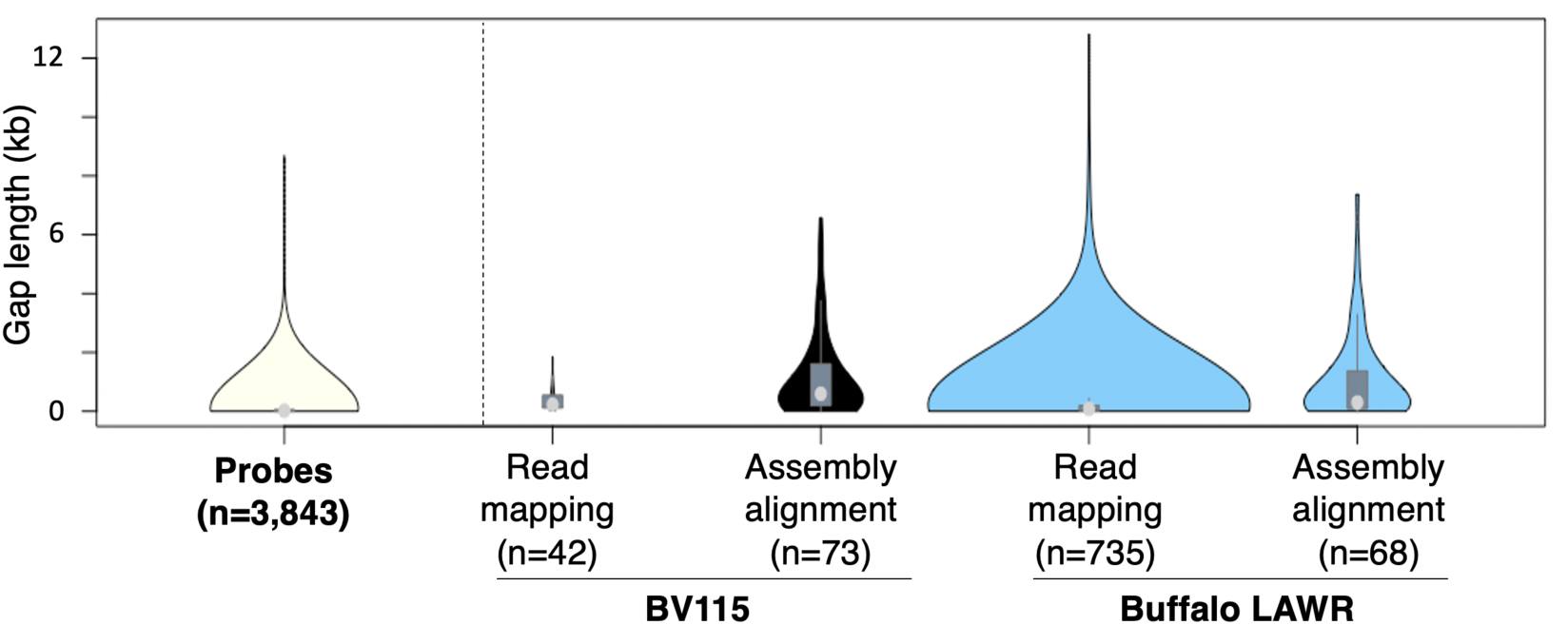
Gaps in coverage of reference genome for capture probes, reads and assemblies. Violin plots of length distribution of gaps in coverage of *T. parva* reference genome by capture probes (n=3,843), as well as for sequence data generated for BV115 (Muguga) and Buffalo LAWR (buffalo-derived) *T. parva* isolates. For each isolate, gaps in coverage were identified after mapping of sequence reads (Read mapping) and after alignment of *de novo* assemblies (Assembly alignment) to the reference *T. parva* genome. For BV115, gaps in coverage after read mapping (n=42) were fewer than observed for Buffalo LAWR (n=735). The number of gaps after alignment of assembly contigs to the reference genome were similar in BV115 (n=73) and Buffalo LAWR (n=68). The Buffalo LAWR sample represents the isolate least similar to the reference and, therefore, to the probes. For the Buffalo LAWR isolate, and in contrast with BV115, assembly alignment resulted in fewer and smaller gaps than read mapping. The median is shown by the light gray circle, and the interquartile range is shown with the dark gray rectangle.

### Data generation

Four *T. parva* isolates were used in this study, which were described in earlier studies (Morzaria et al. 1995; Gotia et al. 2016). Briefly, isolate BV115 consists of infected lymphocytes resulting from the experimental infection of *Bos taurus* animal BV115 with the Muguga reference. This parasite, obtained from the Kenyan highlands, is the source of the reference *T. parva* genome (Gardner et al. 2005), and was also the template for the design of the capture probes. Therefore, enrichment results using the BV115 isolate represent both a proof of principle for this approach and positive control, representing the best possible scenario of a perfect sequence match between probes and the DNA they target. Two other clones derived from cattle infections were used, namely *T. parva* Marikebuni (stabilate 3292), from coastal Kenya, and Uganda (stabilate 3645), from northwest Uganda. These three isolates, originally obtained from cattle, are henceforth designated as “cattle-transmissible” or “cattle-derived”. To determine the success of this approach for buffalo-derived *T. parva*, we used the isolate Buffalo 7014 (stabilate 3081), originally derived from an African Cape buffalo. This is the same isolate for which 36 bp-long whole genome sequence read data was generated by Hayashida and colleagues (Hayashida et al. 2013) and which they referred to as *T. parva* “Buffalo LAWR”; the same nomenclature will be used here, to avoid confusion. Library preparation was initiated using 900 – 1200 ng of total DNA, generated from the extraction of DNA from infected lymphocyte cultures (**Supplemental Table S2**). The proportion of *T. parva* DNA in each sample was 1.9%, 3.1%, 0.9% and 1.7% for BV115, Marikebuni, Uganda and Buffalo LAWR, respectively, with the remainder being host DNA (Gotia et al. 2016) (**Table 1**).

**Table 1.** Specificity and sensitivity of the capture-based parasite DNA enrichment approach.

| Isolate | Pre-enrichment *Theileria* gDNA (%)<sup>a</sup> | Total reads generated<sup>b</sup> | Total reads aligned | Mean coverage<sup>c</sup> | Specificity (%)<sup>d</sup> | Sensitivity (%)<sup>e</sup> |
| --- | --- | --- | --- | --- | --- | --- |
| BV115 | 1.94 | 12,174,316 | 11,952,650 | 146X | 98.03 | 99.80 |
| Marikebuni 3292 | 3.05 | 7,204,556 | 6,740,297 | 202X | 96.47 | 97.70 |
| Uganda 3645 | 0.92 | 5,687,838 | 5,313,295 | 157X | 97.52 | 98.34 |
| Buffalo 7014 | 1.72 | 6,080,972 | 5,446,541 | 160X | 96.40 | 97.59 |

<sup>a</sup> Proportion of the original DNA sample that is composed of *T. parva* DNA, as measured by qPCR (Gotia et al. 2016). <sup>b</sup> Read length for BV115 was 101 bp and 250 bp for the other three strains. <sup>c</sup> Estimation: (Total\_reads\_aligned\*Mean\_read\_length)/genome\_size. The genome size used was the sum of the nuclear and apicoplast reference genomes targeted. <sup>d</sup> Percent reads generated that mapped to *T. parva* reference genome. <sup>e</sup> Percent of the *T. parva* reference genome to which reads map.

For each sample, the gDNA shearing length targeted was 500-700 bp, with average size of captured fragments between 446 and 619 bp. Libraries were sequenced with either Illumina HiSeq 2000 or MiSeq platforms, and 5,687,838 to 12,174,316 sequence reads were generated for each sample and mapped to the *T. parva* reference genome (**Supplemental Table S2**).

### Approach specificity, sensitivity and accuracy

The specificity of the capture approach used is defined here as the fraction of the BV115 sequence reads that map to the parasite reference genome. Specificity of the approach was very high, with >98% of BV115 reads generated mapping uniquely to the *T. parva* reference genome (**Table 1**), a nearly-perfect, 50.5-fold enrichment for parasite DNA. The remaining 1.97% of mapped reads in the present study originated from the host genome.

Sensitivity of the approach is defined here as the fraction of the reference genome to which the BV115 sequence reads map. The sensitivity of this capture approach, based on this probe set, is 99.8%. Not only were all regions of the reference genome against which probes were designed recovered but, in fact, the fraction of the reference genome that is covered by Illumina sequence reads is larger than 95.5%, the fraction of the genome that is covered by probes (**Figure 2, Table 1**). This result demonstrates that we successfully captured segments of the genome that are not included in the probe set, as intended with the experimental design described above. As a result of the high sensitivity of this approach, despite the 3,843 gaps in probe coverage of the *T. parva* genome, the number of segments of the genome with 0X coverage from read mapping was 42, with average length of 402.6 bp (**Figure 1**). This was because repeats or low complexity regions eliminated from the probe set were captured in fragments that also contained neighboring unique regions for which capture by the probes was efficient.

**Figure 2.**
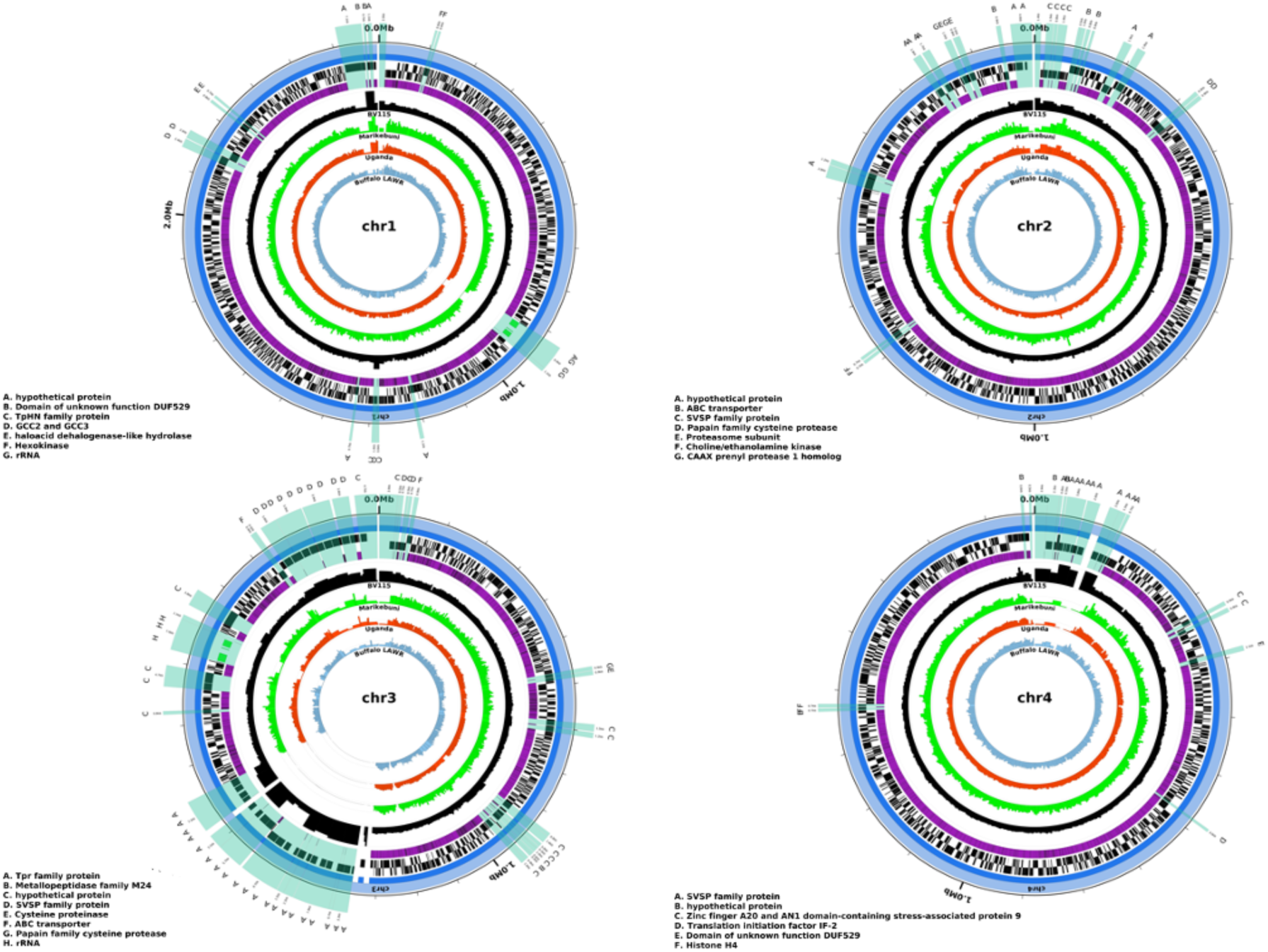
Alignment of probe coverage and read mapping in *T. parva* nuclear chromosomes. For each of the four *T. parva* nuclear chromosomes, starting from the outer-most circle, the following tracks are shown: chromosome scaffolds (light blue); assembly contigs (dark blue); genes encoded in the forward strand, including protein-coding (black), rDNA (green) and tRNA (red) genes; genes encoded in the reverse strand (protein-coding, rDNA, and tRNAs); regions covered by probes (purple); BV115 coverage (absolute read counts in 5kb windows; maximum shown is 500) (black); Marikebuni coverage (green); Uganda coverage (orange); Buffalo LAWR coverage (sky blue). Chromosomal regions without probes that are ≥500 bp are magnified 5000X and highlighted in transparent light green across the tracks showing chromosome, contig, forward and reverse strand genes, and probe coverage. The proteins encoded within these regions without probes are labeled, with the key for each chromosome listed on the bottom left of each plot.

The generation of whole genome sequence (WGS) data based on this capture approach includes one amplification step. To verify the accuracy of the WGS data, and in particular to assess its error rate, we mapped the sequence reads against the reference Muguga genome assembly (Gardner et al. 2005), and identified SNPs. Despite the differences in library protocol, sequencing platforms and data preparation we identified only 107 SNPs across the entire 8.31 Mb genome, for a SNP density of ∼1×10^−5^ SNPs/bp, below the sequencing error of the Illumina platform, and thus providing independent confirmation of the quality of the data generated here (**Supplemental Table S3**).

### Capture specificity and sensitivity for non-reference strains

Both specificity (96.4% - 97.5%) and sensitivity (97.6% - 98.3%) were very high albeit slightly lower for non-reference *T. parva* isolates than they were for BV115 (**Table 1**). However, these values are likely underestimates. Some *T. parva* genes, including those coding for known antigens and some proteins involved in host-parasite interactions, are known to be highly variable, with polymorphism >2%, and possibly much higher (Pelle et al. 2011). This level of polymorphism poses challenges at two levels: *i*) decreased capture efficiency with increasing sequence divergence between probes and target sequences, and (*ii*) lack of read mapping beyond a sequence divergence threshold between sequence reads and genome reference. The former will result in decreased sensitivity while the latter will result in underestimation of both specificity and sensitivity. To determine which of these two factors is responsible for the lower sensitivity in the three non-reference isolates relative to BV115, we generated *de novo* genome assemblies for each isolate, based on the sequence capture data, and compared coverage of each Muguga locus by read mapping to the completeness of the locus sequence extracted from each *de novo* assembly (see section on “Sequence variant detection” below).

### *De novo* genome assemblies based on whole genome sequence capture data

We built several assemblies with the BV115 data, varying the assembly software and genome depth of coverage, and selected the assembly with the longest N50 and cumulative length (not shown). For the reference strain, represented by BV115, nearly every gene was represented in full in the *de novo* genome assembly, and only 7 genes (<0.2% of all genes) were completely absent (**Supplemental Table S4**). The missing genes are located in the most probe-poor regions of the genome, and mainly consist of SVSP multigene family members (Schmuckli-Maurer et al. 2009). In contrast to the assembly data, when the BV115 sequence reads were mapped directly to the reference genome there were no nuclear genes completely or partially missing, demonstrating that despite gaps in probe coverage, we were able to obtain complete nuclear gene coverage in our positive control isolate, and that the fact that some of these genes are missing in the assembly may be due to the difficulty to unambiguously assemble these regions.

The *Tpr* locus, which spans a central region of chromosome 3, is represented in the reference genome assembly by two small contigs plus the edge of one of the two larger chromosome 3 contigs (Gardner et al. 2005). Interestingly, despite the near complete lack of probes in the *Tpr* locus, in BV115 we were able to capture reads that map to most of the locus (**Figure 2**) and reconstruct partially assembled contigs for this chromosomal region (**Figure 3**). This suggests that, in the reference Muguga strain (used to infect animal BV115, and the reference used for probe design), either a *Tpr* gene outside of the *Tpr* locus is similar enough in sequence for probes based on its sequence to cross-capture the *Tpr* locus, or that the few probes present in this *Tpr* locus region were sufficient to anchor and capture sequence fragments that span most of this region of chromosome 3.

**Figure 3.**
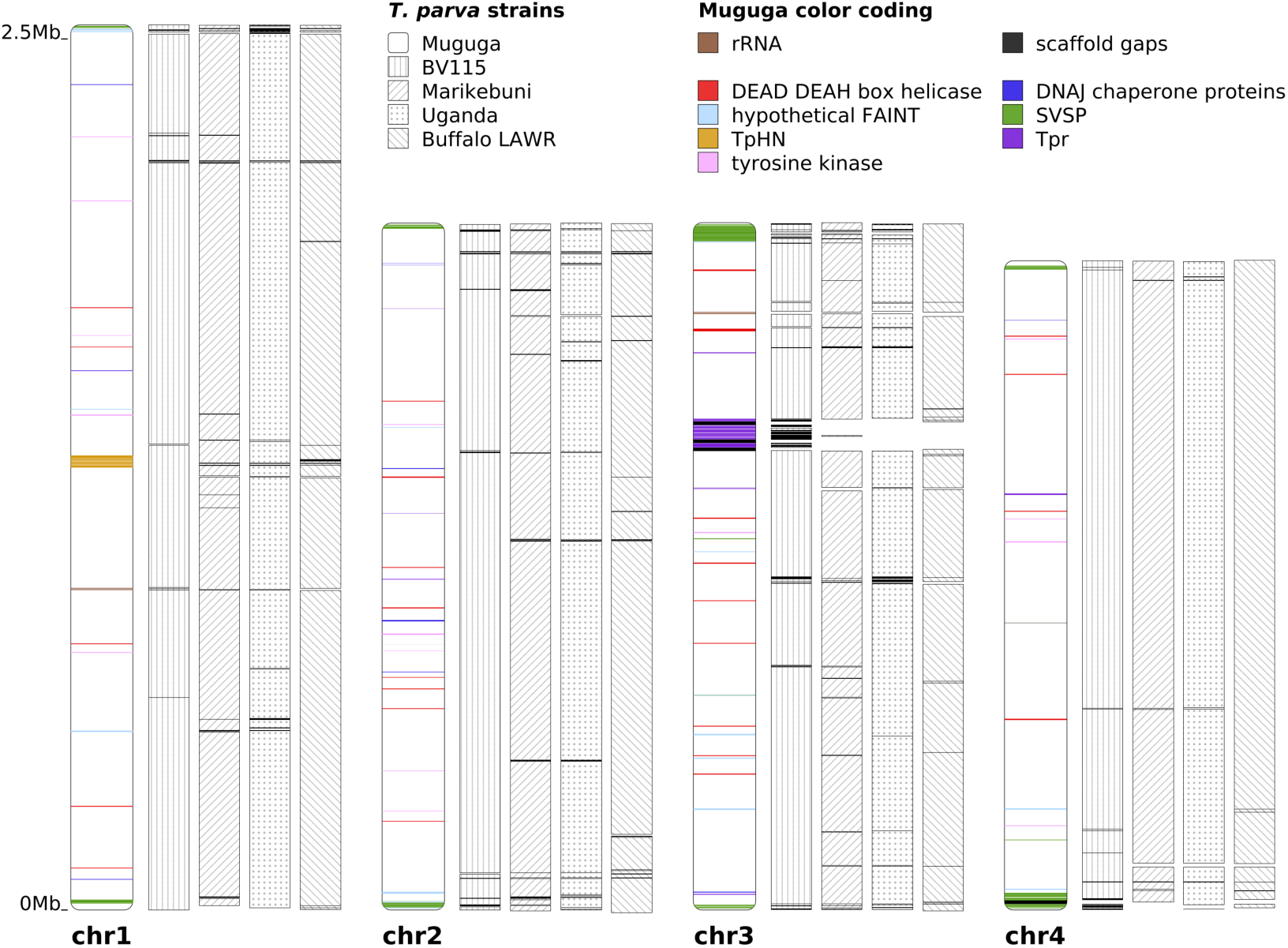
*De novo* genome assemblies for four *T. parva* isolates. In the reference assembly (Muguga), chromosomes 1 and 2 are represented by single scaffolds. The four contigs representing chromosome 3 and the two contigs of chromosome 4 were each merged and are shown as a single scaffold. Assemblies for the four new isolates were generated with SPAdes, and aligned to the *T. parva* Muguga reference, using nucmer. Scaffold placement was centered on the midpoint of each alignment. All gene families with >10 paralogs are colored in the reference Muguga strain. The main genomic regions underrepresented in the new assemblies contain several members of *T. parva* multigene families, including the *Tpr* locus (a total of 24 *Tpr* genes within the two contigs central in chromosome 3), and SVSP family (15 SVSP genes in the first 20kb in the 5’ end of chromosome 4, shown at the bottom).

*De novo* assemblies were also generated for the three non-reference genotypes. They each consist of <130 scaffolds and encompass >95.6% of the reference *T. parva* genome assembly (**Figure 3**; **Supplemental Table S4**). In each case, fewer than 2% of the nuclear genes were completely missing: 56 in Marikebuni, 31 in Uganda, and 27 in Buffalo LAWR. One notable area lacking assembly coverage in all three genome assemblies was the *Tpr* locus. While these genes were not difficult to capture in the BV115, read coverage is poor in the non-reference isolates, suggesting that the reference-based probes failed to capture these genes in other genomes, consistent with the hyper variability of this gene family (Bishop et al. 1997). Of the 37 *Tpr* genes, including dispersed copies present throughout the genome, only six were recovered either partially or completely in all three non-reference genotypes. For each strain, there were several contigs that were not incorporated into the respective genome assembly. These contigs contain sequences with homology to a number of *Theileria parva* genes, a majority of which are members of multigene families or hypothetical proteins (**Supplemental Table S5**).

### Sequence and structural accuracy of *de novo* assemblies

To correct possible errors and validate the *de novo* assemblies, a number of steps were taken. Illumina reads for each of the four strains were mapped to their respective *de novo* assembly. The number of nucleotide differences detected varied between 73 in BV115 and 431 in Buffalo LAWR. Assemblies were then polished with Pilon (Walker et al. 2014) which corrected several of the differences identified (**Supplemental Table S6**). The remainder may represent non-specific read mapping in regions of multigene families, regions with insufficient read coverage for sequence correction or variants segregating in culture at the time of DNA isolation.

Draft assemblies for the non-reference cattle-derived *T. parva* strains used in this study were generated before with 454 data (Henson et al. 2012), which allow validation of the sequence data and assemblies generated here with a capture-based approach. Alignment of the Illumina reads to the 454-based assemblies yielded 203, and 79 SNPs respectively for the Marikebuni and Uganda strains, for a density of <1-2 x 10^−5^ differences per bp. In addition, the BV115 reads were mapped against the Muguga reference genome, with 107 differences identified. All these values fall within the margin of error of Illumina sequencing, again consistent with the high quality of the data generated. Alignment of our *de novo* assemblies to the previously generated 454-based assemblies resulted in a similarly low number of SNPs in each of the three strains (**Supplemental Table S6**).

Finally, when comparing the structure of the BV115 assembly to the Muguga reference, a total of 26 structural variants were detected, with a cumulative length of 11,388 bp (**Supplemental Table S7**). The Marikebuni and Uganda *de novo* assemblies were compared to their respective reference assemblies generated from 454 sequencing data (Henson et al. 2012). The 454-based assemblies consist of 985 and 507 contigs, respectively for Marikebuni and Uganda. These comparisons each yielded 11 or fewer structural variants, totaling at most ∼1000 bp in length (**Supplemental Table S7**). We observed a considerably larger number and cumulative length of structural variants in BV115 relative to its reference compared to what is observed for the other two cattle-derived genotypes, which is possibly an artifact of the high fragmentation of the Marikebuni and Uganda 454 sequencing-based reference genomes (which may prevent the detection of structural differences) relative to the nearly closed Muguga reference. The very low number of cumulative base pairs affected by these variants highlight the high accuracy of the *de novo* assemblies build with the capture data.

### First genome assembly for a buffalo-derived *T. parva* parasite

The generation of *de novo* genome assemblies allows a comprehensive characterization of differences in homologous genomic regions between new strains and the reference. It also makes it possible to characterize missing and unique regions in the new genomes, provided that those regions flank regions that match probes, or that they represent duplicated regions with high sequence similarity to probe-covered genomic segments in the reference. In these situations, *de novo* assemblies also provide a clear advantage for genome characterization over read mapping approaches.

The alignment of the Buffalo LAWR assembly to the Muguga reference revealed >300 structural variants in regions of sequence homology between genomes, affecting a cumulative length of 128,750 bp. These included insertions/expansions totaling a gain of 61,632 bp (83 insertions totaling 17,534 bp in length, 26 tandem duplications totaling 20,000 bp, and 71 repeat expansions totaling 24,098 bp) as well as structural changes resulting in genome reduction, namely deletions and contractions amounting to a loss of 67,118 bp (**Supplemental Table S8**). This suggests that rates of repeat expansion and contraction are fairly balanced.

Overall, however, the genome assembly of the buffalo-derived isolate (Buffalo LAWR) is 8,366,826 bp long (**Supplementary Table S4**), ∼20 Kb longer than the reference *T. parva* Muguga assembly, despite missing genomic regions for which probes were not designed. Compared to that of BV115, which was used as a positive control, the Buffalo LAWR assembly is approximately 130 Kb longer, which supports the hypothesis that the Buffalo LAWR genome is indeed longer. Therefore, we sought to characterize regions unique to the genome of the buffalo parasite.

The automated annotation of the assembly with GeneMark-ES identified new potential genes in regions without reference Muguga gene structures transferred with GMAP to the Buffalo LAWR assembly. In total, 19 genes represented additional copies of *T. parva* genes present elsewhere in the genome, based on top BLAST hits. They encoded mostly hypothetical proteins, plus several putative integral membrane proteins. An additional group of five genes were most similar to homologs found in *Theileria annulata* or *Theileria orientalis* (**Supplemental Table S9**), but not found in *T. parva.* Those with best matches to *T. annulata* included hypothetical proteins, a tRNA-pseudouridine synthase I, and a mitochondrial ribosomal protein S14 precursor gene. The single-copy gene with a best match to *T. orientalis* aligned to a region of chromosome 2 of *T. orientalis* (strain Shintoku) with no genes currently annotated.

These 24 new potential genes described above were run through several prediction software packages for further characterization. There were two predicted proteins with five or greater transmembrane helices, a feature usually associated with transmembrane proteins. There were seven genes predicted to contain signal peptide cleavage sites, and no genes were predicted to be GPI-anchored.

### Sequence variant detection by read mapping and assembly comparison

To determine if the reconstruction of *de novo* draft genome assemblies with the capture data results in additional power to study rapidly evolving antigens compared to the more common read mapping-based approach, we identified sequence variants with both methods. Alignment of sequence reads generated for the two non-reference cattle-derived clones, Marikebuni and Uganda, against the reference genome, followed by stringent SNP calling and filtering, yielded 41,086 and 41,975 SNPs. For the buffalo-derived isolate, Buffalo LAWR, sequence variant calling resulted in 94,999 SNPs (**Supplementary Figure 1**). These values are based on a very high proportion of the genome (**Supplemental Table S3**). These numbers of SNPs are similar to those detected in other cattle-derived isolates in a previous study, which also used a read mapping based approach (Hayashida *et al*. 2013).

A greater number of SNPs was detected by aligning each new assembly to the reference genome and identifying sequence variants (**Supplemental Figure S1**). A total of 55,421 SNPs and 52,385 SNPs were detected in the Marikebuni and Uganda genome assemblies, respectively. Similarly, there were also more SNPs detected in the Buffalo LAWR isolate (n=124,244) based on assembly comparison, than those detected by read mapping, both here and in a previous study using the same strain (Hayashida *et al*. 2013). In each case, ∼25% more SNPs were identified by assembly comparison relative to read mapping.

The discrepancy in SNP counts is due to the fact that reads fail to map to their orthologous regions when the sequence is highly variable between genomes, while these loci are nevertheless present in the *de novo* assembly and can be readily compared across strains. This is further supported by our estimates of gene coverage by each of the methods. In each of the non-reference genotypes, the read mapping approach had, on average, worse gene coverage than the respective coverage using assembly generation (**Figure 4; Supplemental Table S4**). Overall, 98.1% of the nuclear genes were recovered in their entirety in BV115 using both read mapping and assembly alignment. In Marikebuni and Uganda the value was 94.7% and 94.8%, respectively, and 93.6% the Buffalo LAWR strain. Among the 56 nuclear genes not fully recovered in all four genotypes, there were no known single-copy antigens. Most of these genes were located in sub-telomeric regions that are composed of highly repetitive DNA sequences and genes belonging primarily to multigene families (**Figure 3**), while the rest were members of the *Tpr* gene family that occur in the highly repetitive *Tpr* locus of chromosome 3, and hence more difficult to capture since we did not retain probes that mapped to more than five locations in the genome.

**Figure 4.**
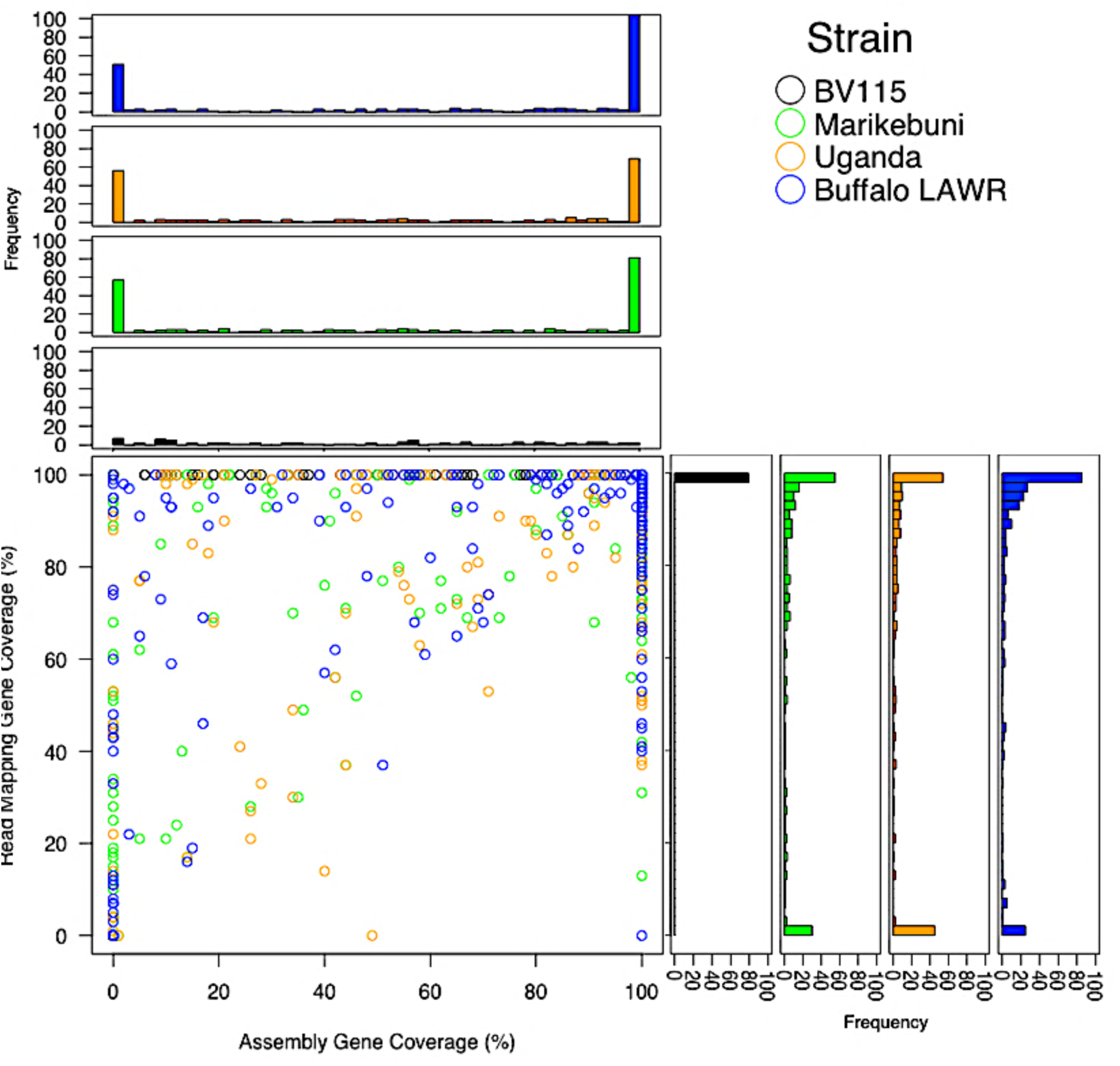
Read mapping and assembly alignment gene coverage in partially covered genes. For each gene of the 4,094 nuclear genes not exhibiting 100% coverage when using both read mapping and assembly alignment to the Muguga reference, the percent of the gene covered using each alignment method is shown in the scatter plot for each of the four isolates. The histograms along the top and right side of the plot show the distribution of gene coverage for read mapping and assembly alignment, respectively. The number of nuclear genes covered 100% using both alignment methods were: BV115 - 4,015; Marikebuni - 3,876; Uganda - 3,883; Buffalo LAWR – 3,831.

### Variation in protein-coding regions

The high degree of completeness of the genome assemblies makes it possible to obtain estimates of the rates of non-synonymous and synonymous polymorphisms per site, *π*_N_ and *π*_S_ respectively, for almost all genes in the genome. This could not be done before due to incomplete read mapping across highly divergent orthologs (Norling et al. 2015). We calculated *π*_N_ and *π*_S_ among the cattle-transmissible isolates (Uganda, Marikebuni and the reference, Muguga), as well as between the Buffalo LAWR isolate and the reference Muguga (**Supplemental Table S10**). Even though estimates of *π*_N_ and *π*_S_ (within species polymorphism) are not reliable estimators of the substitutions rates quantified by *d*_N_ and *d*_S_ (Kryazhimskiy and Plotkin 2008), especially for very small values of *π*, the relative magnitude of *π*_N_ across genes and the ratio *π*_N_/*π*_S_, when defined, may still be informative for the identification of rapidly evolving proteins (Bishop et al. 1997; Jeffares et al. 2007; Weir et al. 2009). Among cattle genotypes, the median *π*_N_ and *π*_S_ across all protein-coding genes were 0.1% and 1.4%, respectively, and the corresponding values for the divergence between a cattle (Muguga) and a buffalo (LAWR) strain were 0.6% and 6.1%, respectively (**Supplemental Table S11)**. As expected, the ratio *π*_N_/*π*_S_ is lower in the Muguga-Buffalo LAWR comparison, since natural selection has had more time to remove slightly deleterious mutations compared to the comparison among cattle strains (**Figure 5a**). Despite the elimination of a relatively higher number of deleterious mutations between Muguga-Buffalo LAWR than among cattle strains, *π*_N_ is still slightly higher in the former comparison because the most recent common ancestor (MRCA) of the Muguga-Buffalo LAWR strains is older than the MRCA of the cattle genotypes (**Figure 5b**). Among 23 well-established antigens and 14 recently identified antigens (Morrison *et al*, in prep.), the average difference in non-synonymous sites between Buffalo LAWR and the ortholog in the reference Muguga was 2.4%, but with a wide range among genes, from 0 to >20%, and a standard deviation of 4.4% (**Table 2; Supplemental Table S12**). Three antigens identified previously as highly polymorphic (Sitt et al. 2018) are the ones with the highest *π*_N_ values between Buffalo LAWR and the reference, namely Tp2, Tp9 and PIM (**Table 2**). Our ability to capture all antigens with near complete coverage using at least one of the approaches demonstrates the value of the capture approach. The ability to fully sequence these antigens in the cattle-and buffalo-derived populations will be useful if future vaccine development against *T. parva* moves toward subunit vaccines (Nene and Morrison 2016).

**Table 2.**
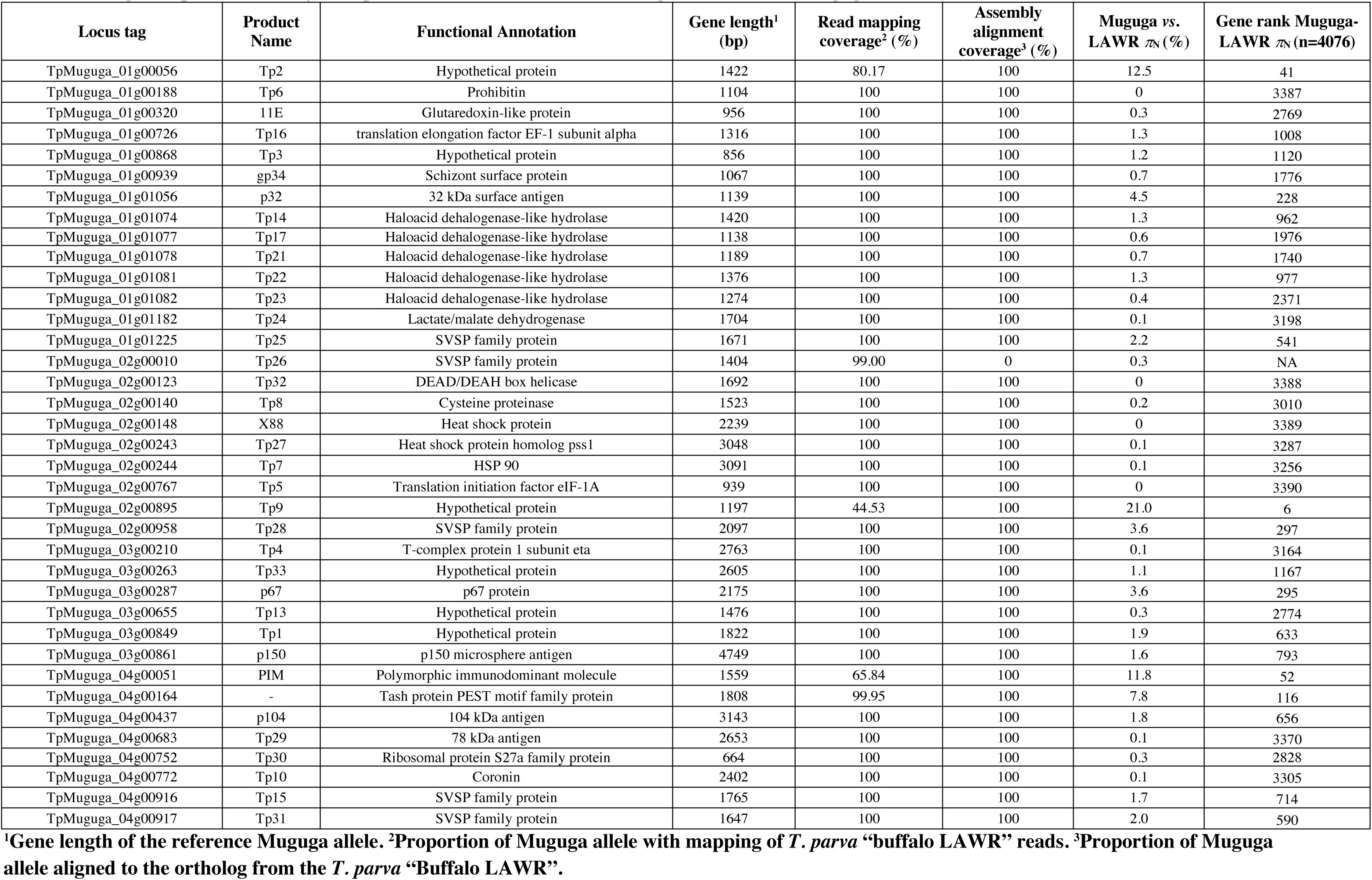
Antigen sequence recovery in *T. parva* “Buffalo LAWR” and divergence relative to Muguga.

| Locus tag | Product Name | Functional Annotation | Gene length<sup>1</sup> (bp) | Read mapping coverage<sup>2</sup> (%) | Assembly alignment coverage<sup>3</sup> (%) | Muguga vs. LAWR $\pi_N$ (%) | Gene rank Muguga-LAWR $\pi_N$ (n=4076) |
| --- | --- | --- | --- | --- | --- | --- | --- |
| TpMuguga_01g00056 | Tp2 | Hypothetical protein | 1422 | 80.17 | 100 | 12.5 | 41 |
| TpMuguga_01g00188 | Tp6 | Prohibitin | 1104 | 100 | 100 | 0 | 3387 |
| TpMuguga_01g00320 | 11E | Glutaredoxin-like protein | 956 | 100 | 100 | 0.3 | 2769 |
| TpMuguga_01g00726 | Tp16 | translation elongation factor EF-1 subunit alpha | 1316 | 100 | 100 | 1.3 | 1008 |
| TpMuguga_01g00868 | Tp3 | Hypothetical protein | 856 | 100 | 100 | 1.2 | 1120 |
| TpMuguga_01g00939 | gp34 | Schizont surface protein | 1067 | 100 | 100 | 0.7 | 1776 |
| TpMuguga_01g01056 | p32 | 32 kDa surface antigen | 1139 | 100 | 100 | 4.5 | 228 |
| TpMuguga_01g01074 | Tp14 | Haloacid dehalogenase-like hydrolase | 1420 | 100 | 100 | 1.3 | 962 |
| TpMuguga_01g01077 | Tp17 | Haloacid dehalogenase-like hydrolase | 1138 | 100 | 100 | 0.6 | 1976 |
| TpMuguga_01g01078 | Tp21 | Haloacid dehalogenase-like hydrolase | 1189 | 100 | 100 | 0.7 | 1740 |
| TpMuguga_01g01081 | Tp22 | Haloacid dehalogenase-like hydrolase | 1376 | 100 | 100 | 1.3 | 977 |
| TpMuguga_01g01082 | Tp23 | Haloacid dehalogenase-like hydrolase | 1274 | 100 | 100 | 0.4 | 2371 |
| TpMuguga_01g01182 | Tp24 | Lactate/malate dehydrogenase | 1704 | 100 | 100 | 0.1 | 3198 |
| TpMuguga_01g01225 | Tp25 | SVSP family protein | 1671 | 100 | 100 | 2.2 | 541 |
| TpMuguga_02g00010 | Tp26 | SVSP family protein | 1404 | 99.00 | 0 | 0.3 | NA |
| TpMuguga_02g00123 | Tp32 | DEAD/DEAH box helicase | 1692 | 100 | 100 | 0 | 3388 |
| TpMuguga_02g00140 | Tp8 | Cysteine proteinase | 1523 | 100 | 100 | 0.2 | 3010 |
| TpMuguga_02g00148 | X88 | Heat shock protein | 2239 | 100 | 100 | 0 | 3389 |
| TpMuguga_02g00243 | Tp27 | Heat shock protein homolog pss1 | 3048 | 100 | 100 | 0.1 | 3287 |
| TpMuguga_02g00244 | Tp7 | HSP 90 | 3091 | 100 | 100 | 0.1 | 3256 |
| TpMuguga_02g00767 | Tp5 | Translation initiation factor eIF-1A | 939 | 100 | 100 | 0 | 3390 |
| TpMuguga_02g00895 | Tp9 | Hypothetical protein | 1197 | 44.53 | 100 | 21.0 | 6 |
| TpMuguga_02g00958 | Tp28 | SVSP family protein | 2097 | 100 | 100 | 3.6 | 297 |
| TpMuguga_03g00210 | Tp4 | T-complex protein 1 subunit eta | 2763 | 100 | 100 | 0.1 | 3164 |
| TpMuguga_03g00263 | Tp33 | Hypothetical protein | 2605 | 100 | 100 | 1.1 | 1167 |
| TpMuguga_03g00287 | p67 | p67 protein | 2175 | 100 | 100 | 3.6 | 295 |
| TpMuguga_03g00655 | Tp13 | Hypothetical protein | 1476 | 100 | 100 | 0.3 | 2774 |
| TpMuguga_03g00849 | Tp1 | Hypothetical protein | 1822 | 100 | 100 | 1.9 | 633 |
| TpMuguga_03g00861 | p150 | p150 microsphere antigen | 4749 | 100 | 100 | 1.6 | 793 |
| TpMuguga_04g00051 | PIM | Polymorphic immunodominant molecule | 1559 | 65.84 | 100 | 11.8 | 52 |
| TpMuguga_04g00164 | - | Tash protein PEST motif family protein | 1808 | 99.95 | 100 | 7.8 | 116 |
| TpMuguga_04g00437 | p104 | 104 kDa antigen | 3143 | 100 | 100 | 1.8 | 656 |
| TpMuguga_04g00683 | Tp29 | 78 kDa antigen | 2653 | 100 | 100 | 0.1 | 3370 |
| TpMuguga_04g00752 | Tp30 | Ribosomal protein S27a family protein | 664 | 100 | 100 | 0.3 | 2828 |
| TpMuguga_04g00772 | Tp10 | Coronin | 2402 | 100 | 100 | 0.1 | 3305 |
| TpMuguga_04g00916 | Tp15 | SVSP family protein | 1765 | 100 | 100 | 1.7 | 714 |
| TpMuguga_04g00917 | Tp31 | SVSP family protein | 1647 | 100 | 100 | 2.0 | 590 |

<sup>1</sup>Gene length of the reference Muguga allele. <sup>2</sup>Proportion of Muguga allele with mapping of *T. parva* “buffalo LAWR” reads. <sup>3</sup>Proportion of Muguga allele aligned to the ortholog from the *T. parva* “Buffalo LAWR”.

**Figure 5.**
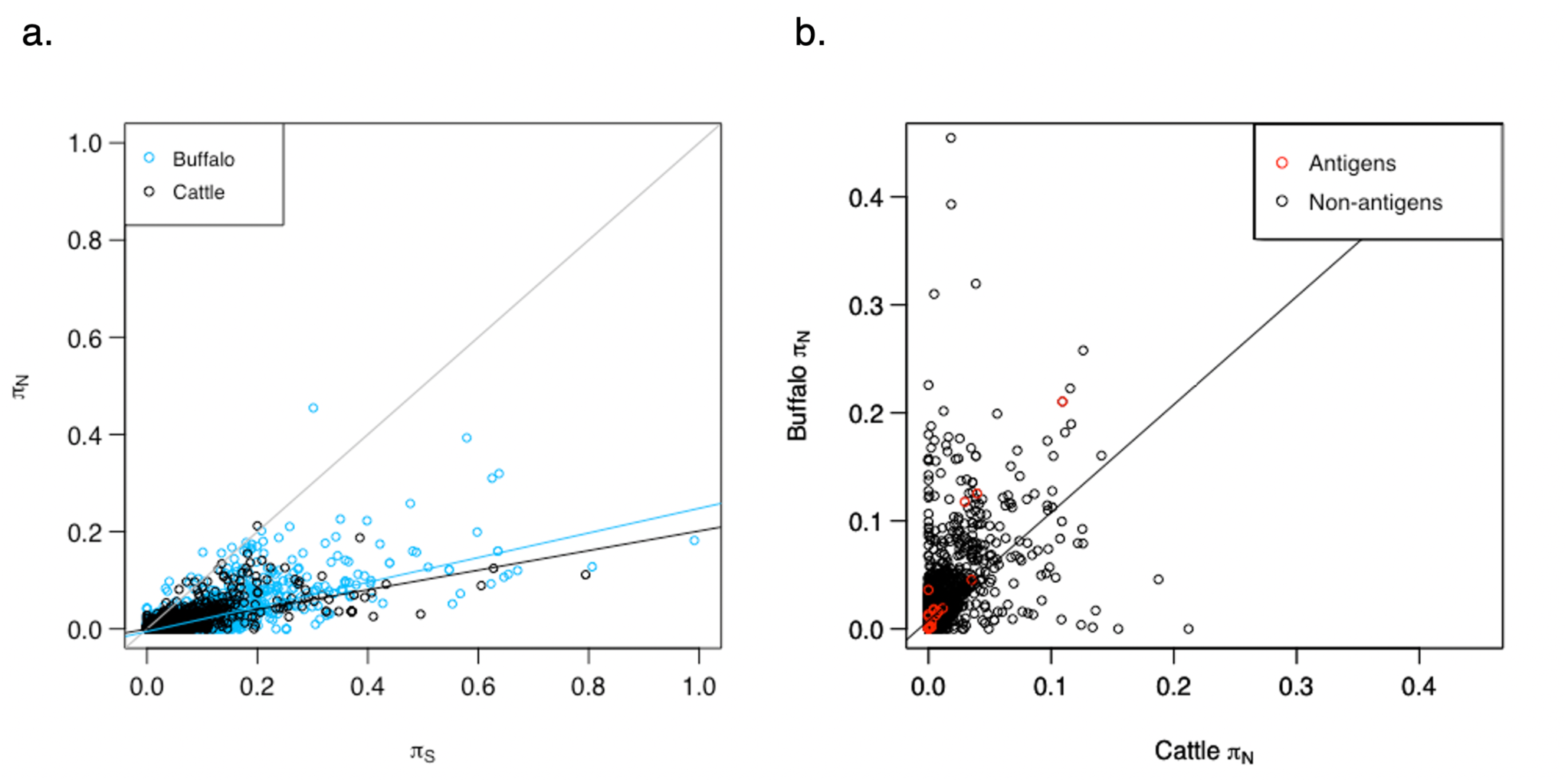
Nucleotide diversity in amino acid-changing (*π*_N_) and silent (*π*_S_) sites. **a.** Rates of nucleotide diversity in non-synonymous (*π*_N_) and synonymous (*π*_S_) sites for all genes in the Muguga reference genome, both among the cattle strains (Muguga, Marikebuni and Uganda), and between the reference Muguga and the Buffalo LAWR. The black regression line corresponds to *π* value for genes among cattle, the light blue regression line corresponds to the *π* value for genes among buffalo, and the black line represents y=x. **b.** For each gene, *π*_N_ among cattle strains (X axis) is compared with *π*_N_ in a buffalo-derived strain relative to the reference Muguga, (Y axis). Known antigens are colored red. The estimated slope regression line is shown, with a slope of 0.99. The R-squared value for the regression is 0.282. Note the shorter axes in 5b.

To identify rapidly evolving genes, we looked for patterns among the 200 (∼5%) genes in three classes: *i*) the highest *π*_N_ value among cattle genotypes, (*ii*) the highest *π*_N_ value between cattle (Muguga) and buffalo (LAWR) genotypes, or (*iii*) the highest *π*_N_/*π*_S_ between cattle (Muguga) and buffalo (LAWR), the most divergent strains. In all three classes, ∼90 genes were annotated as hypothetical, 53 of which were present in two classes and 16 were present in all three classes (**Supplemental Table S13**).

Gene families that were most variable in all three classes included SVSP family proteins, Tpr family proteins, and genes annotated as putative integral membrane family proteins. It is not surprising that the SVSP family genes would have high *π* values, regardless of host, given their localization in sub-telomeric regions and high level of repetitiveness, which are often correlated with higher levels of sequence variation (Schmuckli-Maurer et al. 2009). Likewise, the *Tpr* genes have high levels of sequence variation; this is consistent with their classification as rapidly evolving, antigenic proteins (Bishop et al. 1997; Weir et al. 2010) and, to an extent, to host adaptation (Palmateer *et al*., in prep.). A number of previously identified antigens also appeared on the list, including Tp1 (Graham et al. 2006), which demonstrated a high *π*_N_/*π*_S_ ratio, and Tp2 and Tp9, which appeared in all three classes. These results are consistent with recent findings which demonstrated that, of ten previously studied *T. parva* antigens, these three antigens were observed to have the highest level of nucleotide diversity (Hemmink et al. 2018).

When comparing only *π*_N_ of genes among cattle to *π*_N_ of genes relative to buffalo, the estimated slope was 0.99, showing that the relative rate of non-synonymous divergence across genes is similar in the two populations (**Figure 5b**). However, an *r*^2^ value of 0.282 shows that the regression is not very predictive. Some genes deviate considerably from the regression line, a pattern consistent with differential, host-specific selection in cattle *vs*. buffalo (**Figure 5b**), possibly warranting functional investigation. As expected, known antigens are among the most rapidly evolving genes.

### Genomic divergence between cattle- and buffalo-derived *T. parva*

Hayashida and colleagues (2013) demonstrated that a greater sequence divergence existed between buffalo-derived genotypes and Muguga (103,880-121,545 SNPs), compared to what was observed among cattle-derived isolates (34,814-51,790 SNPs). The same study found that there was no significant evidence for recombination between cattle- and buffalo-derived *T. parva*, the two populations perhaps having evolved a genetic barrier to recombination. Such a barrier may be due to the absence of piroplasm stages in cattle infected with buffalo-derived genotypes and hence the lack of opportunity for co-transmission of cattle- and buffalo-derived parasites in the same tick (Morrison et al. 2015).

To determine genetic differentiation between cattle- and buffalo-derived *T. parva*, and potentially unusually differentiated loci associated with host adaptation, we estimated Wright’s *F*_ST_, which estimates the amount of genetic variation in the population that is due to differences between the two subgroups. *F*_ST_ varies between 0 (panmictic population) and 1 (complete differentiation, with the two subgroups fixed for different alleles at all variable genomic sites). We used the data generated here together with data for distinct strains from Hayashida et al. (2013). This analysis showed a mean genome-wide *F*_ST_ value of 0.436 (**Supplemental Figure S2**). The distribution of *F*_ST_ varies considerably across each of the four nuclear chromosomes, with some regions in the genome nearly fixed for different variants and others homogeneous among cattle- and buffalo-derived strains (**Supplemental Figure S3**). Approximately 3,000 sites (0.036% of the genome), distributed throughout the genome, were found to have an *F*_ST_ value that reached genome-wide significance. These patterns of divergence are consistent with an evolutionary old divergence and extensive population differentiation, preventing the identification of specific genes involved in host-specific strain adaptation. Interestingly, though, this extent of divergence in allele frequency does lend support to the assertion that these are two distinct, host-associated parasite populations. An *F*_ST_ analysis comparing isolates from different studies, collected from the same host, resulted in an *F*_ST_ value of 0 (not shown), ensuring that our comparison between host-associated genotypes is not due to methodological biases of data collected in different studies. These conclusions are somewhat limited by the relatively low genome sequence coverage for samples collected in previous studies (Hayashida *et al.* 2013). A more conclusive study on this topic will require higher coverage from multiple cattle and buffalo *T. parva* genotypes. It is worth noting that previous studies using variable number tandem repeats suggested that *T. parva* populations in co-grazing cattle and buffalo, in central Uganda, were essentially distinct (Oura et al. 2011).

### Apicoplast genomes

The apicoplast genome from the Muguga reference strain is 39,579 base pairs long and contains 70 genes. The probe set covered 64 of the genes either fully or partially (**Supplemental Table S1**). Aligning sequence reads of all three non-reference isolates achieved complete coverage of all 70 genes encoded by the Muguga apicoplast genome. The assemblies also contained the nearly full gene complement of the Muguga apicoplast, as all 70 genes were covered completely in Marikebuni, and 68 were covered completely in the Uganda and Buffalo LAWR apicoplast assemblies. The two partially covered genes in the Uganda strain apicoplast are TpMuguga_05g00034 (37%) and TpMuguga_05g00040 (93%). In the apicoplast of Buffalo LAWR, TpMuguga_05g00037 was covered partially (66%) and TpMuguga_05g00036 was not covered at all.

This fairly complete apicoplast gene set allowed the evaluation of nucleotide diversity per gene. Among cattle isolates, the mean *π*_N_ and *π*_S_ values were both 0.2% **(Supplemental Table S11)**, less than half the mean values for any of the four nuclear chromosomes. The *π*_N_ value between the buffalo isolate and the reference was 0.4%, and the *π*_S_ value was 2.1% (**Supplemental Table S10)**. As expected, these values are greater than the nucleotide diversity among cattle strains, but still considerably lower than the mean nucleotide diversity in nuclear genes among buffalo strains. This suggests that apicoplast genes evolve slower than nuclear genes. Given the role of the apicoplast genes in basic metabolic processes (Aboulaila et al. 2012), they are expected to be highly conserved. Indeed, high levels of apicoplast sequence conservation have been observed across several Apicomplexa species (Huang et al. 2015).

## Discussion

This study demonstrates that high quality whole genome sequence data can be generated for intracellular protozoan-infected mammalian cell lines, despite the large excess of host DNA, using a custom oligonucleotide genome capture array. Additional sequences of *T. parva* have been published since the original *T. parva* reference genome was published in 2005 (Gardner et al. 2005), including those using pyrosequencing technology on DNA extracted from piroplasm-rich red blood cells (Henson et al. 2012), or from lysis of infected lymphocytes (Hayashida et al. 2013). However, the first approach is not feasible on a large scale and the second has low DNA yield and high proportions of host DNA, greatly increasing sequencing costs.

The whole genome capture method described in the present study, with close to 100% specificity and sensitivity, also represents an advancement in targeted enrichment when compared to approaches previously applied to apicomplexan parasites. An implementation of hybrid selection using whole genome “baits” for the parasite *Plasmodium falciparum* yielded an average of 37-fold enrichment with unamplified samples, and no samples with >50% parasite DNA (Melnikov et al. 2011). The application of DNA capture to the apicomplexan *Plasmodium vivax* had a maximum specificity of 80% and sensitivity as low as 84.7% (Bright et al. 2012). Three key differences between the present and previous studies are (*i*) the length of the sheared DNA (here >450 bp, but for example, only 200 bp in (Bright et al. 2012)), (*ii*) the small probe length (here ∼76 bp, compared to 140 bp in Melnikov et al. 2011), and (*iii*) inclusion of probes that map both coding and non-coding regions (e.g., the study by Melnikov and colleagues (2011) included only probes to exons). The first point was intended to facilitate the capture of rapidly evolving or highly variable genomic segments flanking those more conserved and, consequently, matching the probes. The second aimed to maximize the proportion of genome with probe hybridization. Finally, the inclusion of probes to exons as well as introns and intergenic regions was intended to maximize genome recovery. Extreme nucleotide composition, such as close to 100% AT content in some intergenic regions and introns in *P. falciparum*, will limit the applicability of these strategies.

The sequences reported here represent the first whole genome datasets from *T. parva* with sufficient quality and depth of coverage to allow the generation of *de novo* genome assemblies from DNA extracted from infected lymphocyte cultures. This opens a new area for high throughput genotyping of *T. parva* field isolates of both cattle and buffalo origin, and potentially those isolated from the tick vector. As we have shown, *de novo* genome assembly allows for the in-depth characterization of genetic polymorphism, which will provide novel insights into the evolution and population genetics of this parasite and enable the study of rapidly evolving proteins and protein families of interest.

Our study reveals tremendous genetic polymorphism between *T. parva* genotypes, even among just those that are cattle-derived. The average nucleotide diversity among cattle-derived *T. parva* (∼6.5 SNPs/kb) is higher than the 4.23 to 6.29 SNP/kb reported before (Hayashida et al. 2013). This is likely attributed to the high sensitivity and specificity of the approach used here, and the resulting ability to reconstruct fairly complete draft genome assemblies. The SNP density observed is also considerably higher than that seen among strains of the malaria-causing apicomplexan *P. falciparum*, of ∼1 to 2.3 SNP/kb (Miles et al. 2016; Moser et al. 2020). This result is explained by the long co-evolution of *T. parva* with the African Cape buffalo, its asymptomatic carrier, dating back millions of years (McKeever 2009). Therefore, its most recent common ancestor is much older than that of *P. falciparum*, which likely emerged as a result of a relatively recent host transfer from gorilla (Liu et al. 2010). The existence of a broadly effective ECF vaccine, despite the greater SNP density among cattle-derived *T. parva* strains than observed among *P. falciparum* or even *P. vivax* strains, is highly encouraging, as it suggests that *Plasmodium* genetic diversity *per se* is not an insurmountable obstacle to the development of an effective vaccine.

The generation of the first genome assembly for a buffalo-derived *T. parva* strain allows us to address, with a high degree of certainty, several long-standing questions in the field. For example, previous studies based both on discrete loci and low coverage genome-wide data (Pelle et al. 2011; Hayashida et al. 2013), suggested that cattle-derived *T. parva* is significantly less diverse than the buffalo-derived *T. parva* population, an assertion supported by our study, which is based on whole-genome sequence data with very high depth of coverage. We also determined that genome size variation exists between cattle- and buffalo-derived parasites, and identified novel genes in the genome of a buffalo-derived strain, which account for a large proportion of its longer overall genome assembly. It is also noteworthy that the majority of the open reading frames unique to Buffalo LAWR were additional copies of genes present in the *T. parva* reference genome. At this point it remains unknown if this is simply due to the whole-genome capture approach used, which is limited to the probes based on the reference genome and their flanking regions, or if in fact *T. parva* (and eukaryotic parasite genomes in general) have a fairly closed pan genome, in which the acquisition of novel genes is very rare and new gene coding sequences (CDSs) are, instead, primarily the outcome of gene duplications and other gene family expansions, followed by rapid sequence divergence (DeBarry and Kissinger 2011; Kissinger and DeBarry 2011). A larger sample size of genomes analyzed will facilitate the exploration of this issue and elucidate whether the observed pattern is due to gene duplication in buffalo-derived *T. parva* or gene loss in cattle-derived strains.

Characterization of buffalo-derived parasites was necessary to identify differences between the two subsets of parasites. The genome-wide *F*_ST_ value of 0.436 reveals strong differentiation between the two subpopulations of *T. parva*. Despite a small sample size, the resulting *F*_ST_ value is much higher than observed between *P. falciparum* populations sampled across Africa, which ranged between 0.01 – 0.11 (Amambua-Ngwa et al. 2019). We also identified a high SNP density between the buffalo isolate and the Muguga reference - nearly double the SNPs detected among cattle isolates, as well as a large number of structural variants. Adding to this genetic evidence of separate host-associated populations are several epidemiological clues (reviewed in Morrison et al. 2020). These include the fact that tick transmission of buffalo-derived *T. parva* to other cattle has only been achieved on a few occasions and at low efficiency (Neitz 1957; Maritim et al. 1992), and previously described immunological observations that differentiate the two parasites (Radley et al. 1979; Radley 1981). The description of new Apicomplexa species based on genetic information alone is controversial (Krief et al. 2010; Rayner et al. 2011) but not unprecedented (Liu et al. 2017), and it has been proposed that species identification based only on DNA characterization would be more efficient, even though inclusion of other data, such as host and geography, when available, is advisable (Cook et al. 2010). Given the mounting evidence of genomic, immunological and epidemiological differences between cattle- and buffalo-derived *T. parva*, we posit that it is appropriate to return to the original classification of these two parasite populations as separate subspecies (Uilenberg 1981; Norval et al. 1991), or perhaps even discuss their classification as separate species.

A reference genome for a buffalo-derived *T. parva* parasite will allow a more accurate characterization of genetic variation among buffalo-derived strains, in particular in genomic regions that are highly divergent from cattle-derived strains. The generation of multiple buffalo-derived genomes will reveal the proportion of the genome that is variable among these strains, relative to differences that are fixed relative to cattle-derived strains. The potential applications of the capture approach on *T. parva* samples are many and will be valuable in answering translational questions, including improving vaccine design and understanding breakthrough infection by buffalo-derived genotypes in vaccinated cattle (Sitt et al. 2015). Application of the capture approach can also be used to understand the role of heterologous reactivity, or infection with multiple *Theileria* species, that has been implicated as a determinant of the impact of disease control measures at the population level (Woolhouse et al. 2015). Finally, the availability of multiple genome sequences may shed light on the mechanism and frequency of host switching from buffalo to cattle that led to the establishment of the two distinct parasite populations described in this study.

Vaccination remains the most cost-effective tool for prevention of livestock infections and concomitant cattle morbidity and mortality. Apicomplexan parasites of livestock are often closely related to human-infective species with respect to the protective immune responses induced, and therefore represent potential models for evaluation of responses to human infection (McAllister 2014). However, while there are several veterinary vaccines against protozoa that have been manufactured by veterinary authorities in collaboration with the private sector for decades, there is still no fully efficacious vaccine against any protozoan parasites that infect humans (Meeusen et al. 2007). A primary challenge to the development of such vaccines is the high degree of antigenic polymorphism, making the collection of this information critical (Takala and Plowe 2009; Barry and Arnott 2014). The problem is particularly acute when the pathogen parasitizes nucleated host cells, resulting in an extremely small ratio of host-to-parasite DNA (Gotia et al. 2016). This presents a substantial obstacle for species in the genus *Theileria*, as well as for bacterial pathogens such as *Chlamydia* and *Rickettsia* (Bachmann et al. 2014; Merhej et al. 2014; Gotia et al. 2016). The approach described here offers a major advance in the capacity to characterize genetic diversity of intracellular protozoan parasite populations, which potentially can enhance informed development of more broadly efficacious vaccines, including protective vaccines against buffalo-derived *T. parva*. To achieve the widest protection, any future development of subunit vaccines against *T. parva* should consider the inclusion of orthologs from buffalo-derived strains.

## Methods

### Samples and parasite-host ratio

Four *T. p*arva isolates, described originally in Morzaria *et al*. (1995) were used. Schizont-infected bovine lymphocyte cultures were derived from lymph node biopsies taken from cattle experimentally infected with *T. parva* sporozoite stabilates Marikebuni_3292, Uganda_3645, Muguga_3087 and Buffalo_7014_3081. The Uganda and Marikebuni stabilates were clonal (Morzaria et al. 1995), whereas the Muguga and Buffalo_7014 stabilates were not cloned and therefore potentially contained multiple parasite genotypes. The Buffalo_7014 isolate was analyzed as Buffalo LAWR by Hayashida and colleagues (Hayashida et al. 2013). Bovine lymphocytes infected with the schizont stage of each isolate were propagated using established protocols (Maramorosch 2012). DNA was extracted from schizont-infected lymphocyte cell line cultures using standard protocols (Sambrook et al. 1989), including proteinase K digestion, phenol/chloroform extraction and ethanol precipitation. The ratio of parasite to host DNA was estimated for each sample, using a qPCR-based approach to estimate the absolute DNA amount separately for bovine and *T. parva* DNA (Gotia et al. 2016).

### Genomic library construction

Library preparation was initiated using 900 – 1200 ng of total DNA, generated from the extraction of total DNA from infected lymphocyte cultures (**Supplemental Table S2**). Paired-end (PE) genomic DNA libraries were constructed for sequencing on Illumina platforms using the NEBNext® DNA Sample Prep Master Mix Set 1 (New England Biolabs, Ipswich, MA). First, DNA was sheared with the Covaris E210, to fragments targeted to 500-700 bp in length. Then libraries were prepared using a modified version of manufacturer’s protocol. The DNA was purified between enzymatic reactions and the size selection of the library was performed with AMPure XT beads (Beckman Coulter Genomics, Danvers, MA).

### Whole-genome DNA sequence capture

A custom-designed Nimblegen SeqCap EZ oligo library was used to target capture the *T. parva* genomic DNA in each genomic library for high-throughput sequencing using the 454 GS FLX and Illumina HiSeq2000 and MiSeq platforms. The capture method utilizes custom-designed, biotinylated oligonucleotides for hybridization to the target sequence. The custom oligo library used here was designed based on the *T. parva* Muguga reference genome sequence with accession number AAGK01000000. Following hybridization of library fragments to the oligo baits, streptavidin-coated magnetic beads are used to capture the bound fragments, and unbound fragments are washed away leaving captured library fragments ready for sequencing.

### Sequencing

The BV115 Illumina PE library was sequenced using the 100 bp paired-end protocol on an Illumina HiSeq2000 sequencer, using approximately 7.25% of a flowcell lane. The libraries for the remaining three isolates were sequenced on a MiSeq platform, using the 250 bp paired-end protocol, multiplexed into a single run, with each using roughly 1/3 of the sequencing capacity. Raw data from the sequencers was processed using Illumina’s RTA and CASAVA pipeline software, which includes image analysis, base calling, sequence quality scoring, and index demultiplexing. Data was then processed through both FastQC (http://www.bioinformatics.bbsrc.ac.uk/projects/fastqc/) and in-house pipelines for sequence assessment and quality control. These pipelines report numerous quality metrics and perform a megablast-based contamination screen. By default, our quality control pipeline assesses base call quality and truncates reads where the median Phred-like quality score falls below Q20.

### Read mapping and genome assembly

The Illumina sequence data were aligned to the reference genome (accession number AAGK01000000) using the Bowtie 2 Aligner (Langmead and Salzberg 2012). Statistics for depth of coverage (number of reads mapped per position) and reference genome breadth of coverage (fraction of the reference genome to which reads map) were generated using internal protocols. Gene coverage was calculated using genomecov from the bedtools suite of tools (Quinlan and Hall 2010). The Illumina data were assembled using the SPAdes Assembler v3.9.0 (Nurk et al. 2013). Because of coverage limitations inherent in almost all genome assembler software, the high-coverage Illumina data was randomly sub-sampled to depths of coverage ranging between 10X and 200X in 10X and 25X increments. The optimal assembly in each case was selected using a combination of statistics including total contig count, contig N50 (contig length for which the set of all contigs of that length or longer contains at least half of the assembly), maximum contig length and total assembly length. The optimal assembly for the BV115 and Uganda assemblies had read coverage cutoff values of 25X, and the optimal Marikebuni and Buffalo LAWR assemblies had read coverage cutoff values of 10X. Assembly contigs were evaluated for host contamination and any contigs matching to the host were removed. In-house scripts were used to determine the extent of overlap between new assemblies and the reference genome (reference genome breadth of coverage) and to generate statistics regarding genes present or absent from the new assemblies. Assembly correction was done using Pilon (v1.22) (Walker et al. 2014), using default parameters, with the respective Illumina sequencing reads for each strain. Assembly gene coverage for each strain was calculated using in-house scripts (https://github.com/jorvis/biocode/blob/master/general/calculate_gene_coverage_from_assembly.py).

### Single nucleotide polymorphism (SNP) and structural variant detection and characterization

The Bowtie 2 alignments were converted to a BAM file using SAMtools (Li et al. 2009). The Genome Analysis Toolkit (GATK) (McKenna et al. 2010) was used to identify and correct misalignments caused by small indels, and then to call both SNPs and indels. The resulting VCF file was filtered with stringent criteria to eliminate potentially false SNPs, requiring depth greater than 12, quality greater than 50, phred-scaled p-value using Fisher’s Exact Test less than 14.5, and Root Mean Square mapping quality zero less than 2. The SNPs were classified by location into intergenic, intronic, synonymous, non-synonymous, read-through or non-sense using VCFannotator (vcfannotator.sourceforge.net). SNPs were detected in assemblies using the show-snps option of the MUMmer3 (v3.23) (Kurtz et al. 2004). *F*_ST_ (as implemented by Weir and Cockerham (Weir and Cockerham 1984)) was estimated using VCFtools v0.1.14 (Danecek et al. 2011). The updated *T. parva* Muguga genome annotation (Tretina et al. 2020b) was transferred on to the new assemblies using the Genomic Mapping and Alignment Program (GMAP) v2014-04-06 (Wu and Watanabe 2005). Assemblytics was used to identify structural variants between the *de novo* assemblies and the reference genome or corresponding assemblies generated from 454 sequencing (Nattestad and Schatz 2016).

### Nucleotide diversity estimation

Nucleotide diversity (the average number of nucleotide differences per site, *π*), was calculated based on SNPs called from read mapping and from assembly comparison. Nucleotide diversity from read mapping was done using the VCFtools v0.1.14 package with the --site-pi option (Danecek et al. 2011). This approach requires reads to map across isolates and does not correct for multiple hits. Nucleotide diversity from *de novo* assemblies was estimated for the CDS alignment for each gene, using the Nei-Gojobori method, which corrects for multiple hits (Nei and Gojobori 1986). In our study, this approach requires that the locus be present in the *de novo* assemblies.

### Gene prediction

To identify novel genes in contigs and genomic segments with no gene annotations we used the gene prediction software Genemark-ES (Lomsadze et al. 2005). To ensure that these contigs are truly part of the *T. parva* genome, all new predicted genes were contained within contigs encoding homologs to *T. parva* genes. To identify true genes, we used BLASTN to search each predicted gene model against NCBI’s non-redundant nucleotide database (Camacho et al. 2009), selecting only those with E-value less than 1×10^−5^. Genes matching to multigene family members were removed. Prediction of transmembrane helices in proteins was done using TMHMM2.0 (Sonnhammer et al. 1998). Prediction of GPI-anchor sites was done using PredGPI (Pierleoni et al. 2008). The presence and location of signal peptide cleavage sites in amino acid sequences was predicted using SignalP 4.1 (Nielsen 2017).

## Data Access

Raw sequence reads and corresponding genome assemblies available from NCBI under project number PRJNA16138. (In progress)

## Acknowledgements

This work was supported in part by the by the Bill and Melinda Gates Foundation award OPP1078791 (NCP, KT, OOI, HTG, VN, JCS) and by the United States Department of Agriculture, Agricultural Research Service (USDA-ARS; through agreement #59–5348–4-001 to JCS). This work was also supported in part by an Immunity and Infection T32 training grant, NIH/NIAID T32 AI007540-14 (KT), and by NIH award R01 AI141900 (NCP, JBM, JCS). All sequencing was done at the Institute for Genome Sciences’ Genomics Resource Center.

