## Supplemental Figures and Tables for "Capture-based enrichment of *Theileria parva* DNA enables full genome assembly of first buffalo-derived strain and reveals exceptional intra-specific genetic diversity"

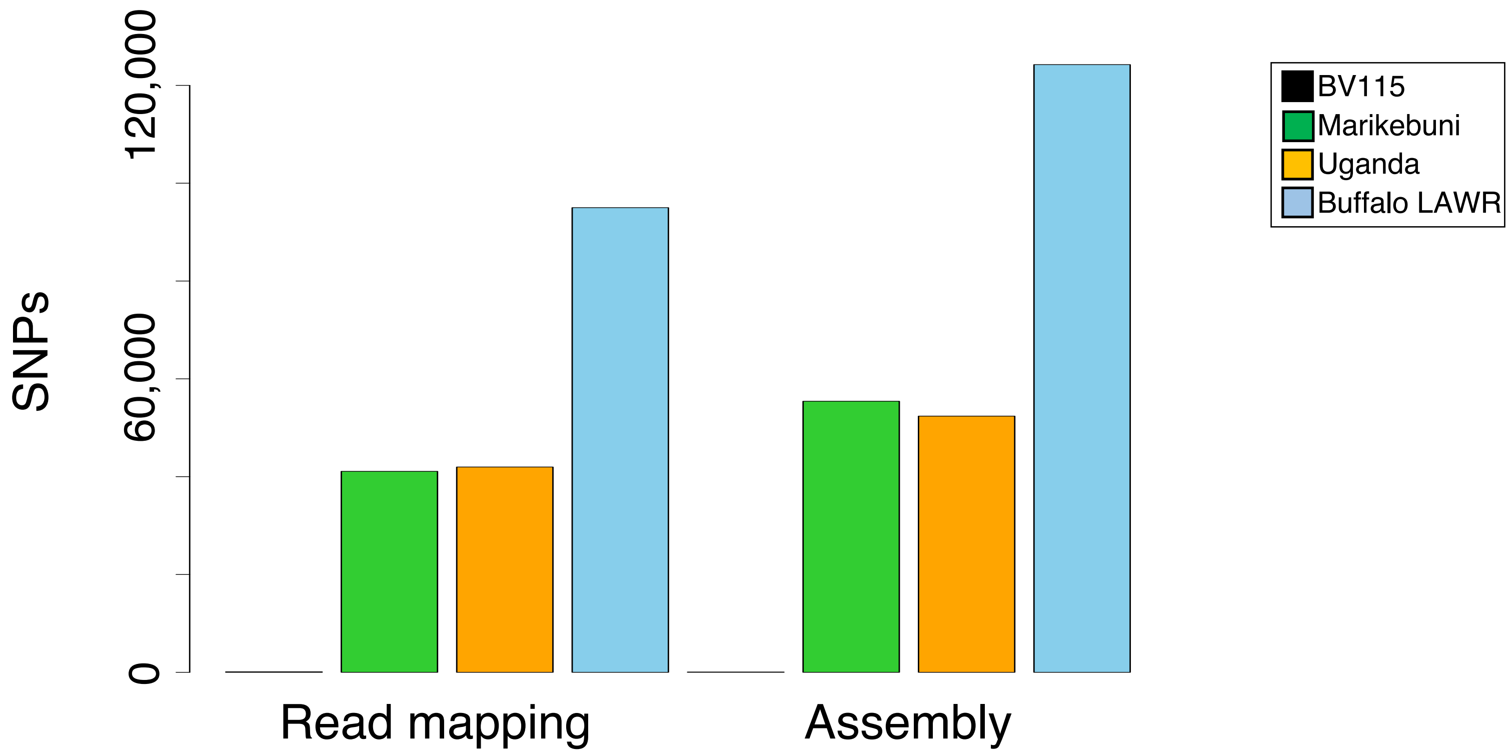


**Supplemental Figure S1. Total count of single nucleotide polymorphisms (SNPs) per sample.** The total number of SNPs identified was compared between read-mapping and assembly approaches, for each of the four isolates. As expected, almost no SNPs were found between BV115 (animal infected with the Muguga strain) and the reference *T. parva* Muguga genome assembly. Identification of SNPs based on assembly alignment is consistently more sensitive than red mapping. Twice as many SNPs are found in the buffalo- than in cattle- derived strain relative to the Muguga reference, from cattle.

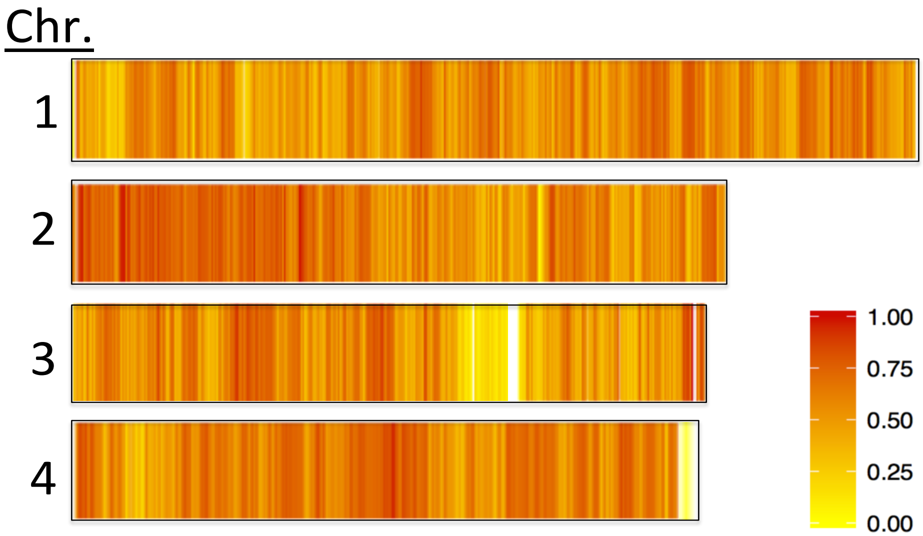


**Supplemental Figure S2. Genetic differentiation between cattle- and buffalo-derived strains.** Window-based $F_{ST}$ analysis comparing cattle strains to those derived from buffalo. Analysis includes our strains and those from Hayashida *et al*. (2013). The windows used were 4,000 bp long with a 1,000 bp overlapping window. Genome-wide $F_{ST}$ was 0.436.

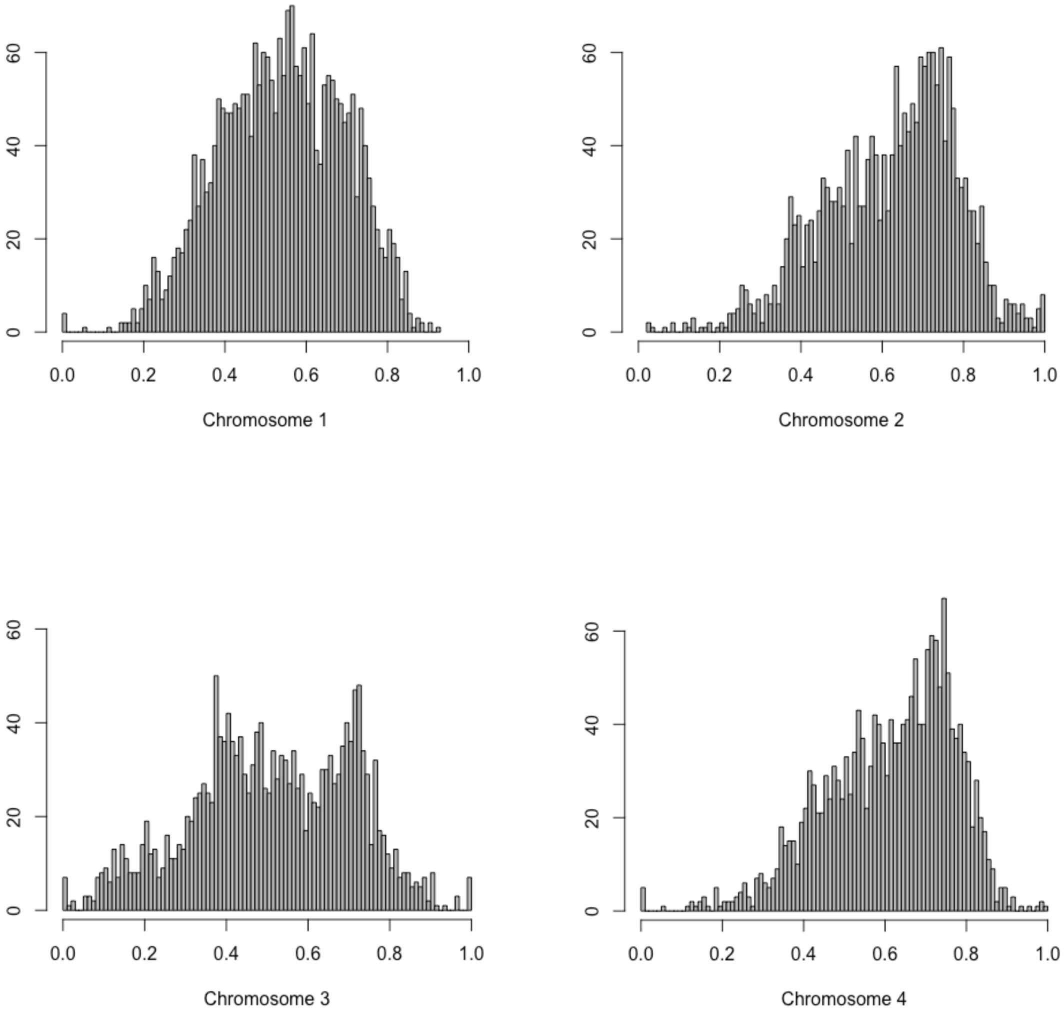


**Supplemental Figure S3. Distribution of $F_{ST}$ values within nuclear chromosomes.** Histograms of window-based $F_{ST}$ values calculated for each nuclear chromosome, showing a wide range of $F_{ST}$ values throughout the genome. Frequency of average $F_{ST}$ value is shown on x-axis, $F_{ST}$ values shown on y-axis.

**Supplemental Table S1. Probe Coverage Statistics.** Properties of gaps in probe coverage across all assembly contigs and supercontigs (chromosomes) and across protein-coding genes.

| Chromosome | Contig | Contig Length (bp) | Genomic Regions Not Mapped by Probes | | | | | | | Gene Mapping by Probes | | | |
|---|---|---|---|---|---|---|---|---|---|---|---|---|---|
| | | | Number of gaps | Median length (bp) | Mean length (bp) | StdDev (bp) | Min length (bp) | Max length (bp) | Cumulative (bp) | Total Genes (#) | Genes Not Covered (#) | Partially Covered Genes (#) | Fully Covered Genes (#) |
| 1 | AAGK01000001 | 2,540,030 | 1,183 | 17 | 63 | 325 | 2 | 7,131 | 74,943 | 1224 | 4 | 570 | 650 |
| 2 | AAGK01000002 | 1,971,884 | 935 | 19 | 81 | 257 | 2 | 4,448 | 76,092 | 984 | 7 | 462 | 515 |
| 3 | AAGK01000005 | 1,317,241 | 755 | 21 | 74 | 206 | 2 | 3,945 | 56,002 | 635 | 0 | 313 | 322 |
| 3 | AAGK01000006 | 570,487 | 329 | 23 | 200 | 866 | 2 | 7,210 | 65,647 | 283 | 14 | 139 | 130 |
| 3 | AAGK01000007 | 41,585 | 22 | 126 | 1,569 | 2,559 | 7 | 8,690 | 34,507 | 18 | 9 | 9 | 0 |
| 3 | AAGK01000008 | 13,275 | 1 | 13,275 | 13,275 | - | 13,275 | 13,275 | 13,275 | 6 | 6 | 0 | 0 |
| 4 | AAGK01000004 | 1,835,834 | 762 | 14 | 46 | 130 | 2 | 2,010 | 35,032 | 934 | 3 | 397 | 534 |
| 4 | AAGK01000003 | 17,691 | 21 | 153 | 670 | 1,083 | 16 | 3,808 | 14,069 | 10 | 4 | 6 | 0 |
| Apicoplast | AAGK01000009 | 39,579 | 29 | 8 | 204 | 490 | 2 | 1,860 | 5,910 | 70 | 6 | 22 | 42 |
| Total | 9 | 8,347,606 | 4,037 | 6 | 87 | 454 | 2 | 52,377 | 375,477 | 4,164 | 53 | 1,918 | 2,193 |

**Supplemental Table S2. Sample metadata.** Properties related to library construction and generation of whole genome sequence data.

| Isolate | Sequencing Platform | Starting Material (ηg) | Average gDNA shearing size (bp) | Average Size of Captured Fragment [a] | Mean Read Length (bp) | Reads generated |
|---|---|---|---|---|---|---|
| BV115 | Illumina HiSeq 2000 | 900 | 500 | 446 | 101 | 12,174,316 |
| Marikebuni | Illumina MiSeq | 900 | 600-700 | 533 | 250 | 7,204,556 |
| Uganda | Illumina MiSeq | 900 | 600-700 | 555 | 247 | 5,687,838 |
| Buffalo LAWR | Illumina MiSeq | 1,200 | 600-700 | 619 | 249 | 6,080,972 |

**Supplemental Table S3. Sequence variants identified by read mapping relative to the reference *T. parva* Muguga genome assembly.**

| | ***T. parva* Muguga genome with coverage that satisfies SNP calling filter** | **Sequence variants *vs*. reference *T. parva* Muguga genome[1]** | |
|---|---|---|---|
| **Isolate** | **Base pairs (%)** | **SNPs** | **INDELs** |
| BV115 | 99.50 | 107 | 18 |
| Marikebuni | 96.81 | 41,086 | 11 |
| Uganda | 97.48 | 41,975 | 7 |
| Buffalo LAWR | 95.61 | 94,999 | 4 |

[1]Sequence variants, including single nucleotide polymorphisms (SNPs) and small insertions and deletions (INDELs) were identified using the Genome Analysis Toolkit.

**Supplemental Table S4. Gene content in *de novo T. parva* genome assemblies.** Assembly length, number of contigs and gene content are shown. Gene content was assessed with two approaches: "read mapping"[1] and "assembly alignment"[2].

| Sample | Assembly Length (bp) | # Scaffolds | Complete Nuclear Genes | | Partial Nuclear Genes | | Absent Nuclear Genes | | *T. parva* Reference Genome Aligned (%) |
|---|---|---|---|---|---|---|---|---|---|
| | | | Assembly Alignment | Read Mapping | Assembly Alignment | Read Mapping | Assembly Alignment | Read Mapping | |
| BV115 | 8,236,259 | 123 | 4,015 | 4,094 | 72 | 0 | 7 | 0 | 98.05 |
| Marikebuni | 8,274,714 | 116 | 4,024 | 3,917 | 84 | 147 | 56 | 30 | 96.18 |
| Uganda | 8,277,618 | 126 | 4,020 | 3,922 | 89 | 128 | 31 | 44 | 96.16 |
| Buffalo LAWR | 8,366,826 | 109 | 4,017 | 3,871 | 96 | 198 | 27 | 25 | 95.68 |

[1]Proportion of each gene in the reference genome assembly with read coverage from each isolate. [2]Proportion of each gene in the reference genome that is aligned with an orthologous section in the de novo assembly of each non-reference isolate. These results were split into three categories: genes with complete coverage, genes partially covered, and genes that are completely absent from the assembly.

**Supplemental Table S5. Homology searches for predicted *Theileria parva* genes in unmapped contigs.**

| | Strain | | | |
|---|---|---|---|---|
| | **BV115** | **Marikebuni** | **Uganda** | **Buffalo LAWR** |
| Total Unmapped contigs | 93 | 61 | 72 | 82 |
| Contigs with no defined orthology to the reference *T. parva* genome | 6 | 9 | 7 | 12 |
| **Gene family** | | | | |
| SVSP family protein | 39 | 24 | 28 | 17 |
| Tpr family protein | 28 | 28 | 0 | 30 |
| TpHN family protein | 14 | 7 | 0 | 9 |
| DEAD/DEAH box helicase | 0 | 0 | 1 | 2 |
| hypothetical protein | 24 | 41 | 64 | 67 |
| ABC transporter | 7 | 12 | 4 | 12 |
| Other | 16 | 15 | 34 | 28 |
| **Total** | 128 | 127 | 131 | 165 |

*Portions of the above gene families were found in the unmapped contigs

**Supplemental Table S6. Assembly validation and correction.**

| **Validation Test** | **BV115** | **Marikebuni** | **Uganda** | **Buffalo LAWR** |
|---|---|---|---|---|
| # SNPs when aligning reads to respective *de novo* assembly | 73 | 320 | 183 | 431 |
| # SNPs in *de* novo assembly corrected with Pilon | 50 | 139 | 78 | 117 |
| # SNPs when aligning reads to respective reference assembly[1] | 107 | 203 | 79 | NA |
| # SNPs when aligning *de novo* assemblies against reference assembly[1] | 92 | 96 | 165 | NA |
| # SNPs when aligning *de novo*, Pilon-corrected assemblies against reference assembly[1] | 61 | 85 | 158 | NA |

[1] References used: for BV115 the reference Muguga; for Marikebuni and Uganda, the 454-based assemblies in Henson *et al*. (2012)

**Supplemental Table S7. Structural variants in *de novo* assemblies compared to reference assemblies[1].**

| | BV115 | | Marikebuni | | Uganda | |
|---|---|---|---|---|---|---|
| **Variant type** | **Count** | **Total bp** | **Count** | **Total bp** | **Count** | **Total bp** |
| **Insertion** | 2 | 171 | 0 | 0 | 0 | 0 |
| **Deletion** | 1 | 128 | 1 | 51 | 0 | 0 |
| **Tandem expansion** | 10 | 5,206 | 9 | 914 | 5 | 373 |
| **Tandem contraction** | 6 | 1,099 | 1 | 87 | 0 | 0 |
| **Repeat expansion** | 1 | 69 | 0 | 0 | 0 | 0 |
| **Repeat contraction** | 6 | 4,715 | 0 | 0 | 0 | 0 |
| **Total for all variants** | 26 | 11,388 | 11 | 1,052 | 5 | 373 |

[1] BV115 assembly was compared to the reference *T. parva* Muguga genome (Gardner et al. 2005). The assemblies for Marikebuni and Uganda were aligned to respective references previously generated with 454 data (Henson et al. 2012). Tandem expansions and contractions occur between overlapping variants, whereas repeat expansions and contractions occur within unmappable gaps between alignments.

**Supplemental Table S8. Structural variants between *T. parva* strains[1].**

| | Marikebuni | | Uganda | | Buffalo LAWR | |
|---|---|---|---|---|---|---|
| **Variant type** | **Count** | **Total bp** | **Count** | **Total bp** | **Count** | **Total bp** |
| **Insertion** | 32 | 2,955 | 26 | 3,363 | 83 | 17,534 |
| **Deletion** | 19 | 1,725 | 25 | 2,332 | 66 | 18,941 |
| **Tandem expansion** | 10 | 10,840 | 13 | 16,340 | 26 | 20,000 |
| **Tandem contraction** | 6 | 6,794 | 6 | 8,819 | 8 | 1,142 |
| **Repeat expansion** | 64 | 25,151 | 67 | 23,080 | 71 | 24,098 |
| **Repeat contraction** | 69 | 31,027 | 60 | 24,394 | 78 | 47,035 |
| **Total for all variants** | 200 | 78,492 | 197 | 78,328 | 333 | 128,750 |

[1] Each assembly was compared to the reference *T. parva* Muguga genome (Gardner et al. 2005).

**Supplemental Table S9. Best BLAST match for each *T. parva* LAWR gene without a detectable homolog in *T. parva* Muguga.**

| Species | Product name | e-value | length |
|---|---|---|---|
| *Theileria annulata* | mitochondrial ribosomal protein S14 precursor | 5.89E-122 | 448 |
| *Theileria annulata* | hypothetical protein | 8.90E-159 | 571 |
| *Theileria annulata* | hypothetical protein | 1.53E-82 | 316 |
| *Theileria annulata* | tRNA-pseudouridine synthase I | 1.39E-168 | 604 |
| *Theileria annulata* | hypothetical protein | 0 | 776 |
| *Theileria orientalis* | Match to chromosome 2 | 8.74E-31 | 145 |

**Supplemental Table S10. Nucleotide diversity among cattle-derived *T. parva* strains (Muguga, Marikebuni and Uganda), and between cattle (Muguga reference) and the buffalo-derived strain *T. parva* Buffalo LAWR.**

See Excel file Supplemental Table S10

**Supplemental Table S11. Summary statistics of nucleotide diversity among cattle-derived *T. parva* strains, and between cattle and the buffalo-derived strain *T. parva* Buffalo LAWR shown in Supplemental Table 10.**

ALL LOCI

| | | Buffalo | | | | | | | | Cattle | | | | | | | |
|---|---|---|---|---|---|---|---|---|---|---|---|---|---|---|---|---|---|
| | | $\pi_N$ | | | | $\pi_S$ | | | | $\pi_N$ | | | | $\pi_S$ | | | |
| | | Mean | Median | St Dev | Max | Mean | Median | St Dev | Max | Mean | Median | St Dev | Max | Mean | Median | St Dev | Max |
| **Nuclear genome** | **(n=4032)** | **0.013** | 0.006 | 0.025 | 0.455 | **0.072** | 0.061 | 0.064 | 1.207 | **0.005** | 0.001 | 0.013 | 0.212 | **0.027** | 0.014 | 0.045 | 0.628 |
| Chrm 1 | (n=1,206) | 0.011 | 0.006 | | 0.393 | 0.072 | 0.064 | | 1.207 | 0.005 | 0.002 | | 0.126 | 0.030 | 0.025 | | 0.628 |
| Chrm 2 | (n=974) | 0.011 | 0.004 | | 0.258 | 0.060 | 0.050 | | 0.654 | 0.005 | 0.000 | | 0.155 | 0.021 | 0.005 | | 0.433 |
| Chrm 3 | (n=925) | 0.019 | 0.008 | | 0.455 | 0.093 | 0.071 | | 0.646 | 0.008 | 0.002 | | 0.212 | 0.041 | 0.024 | | 0.496 |
| Chrm 4 | (n=927) | 0.011 | 0.005 | | 0.178 | 0.066 | 0.061 | | 0.806 | 0.004 | 0.000 | | 0.187 | 0.017 | 0.003 | | 0.385 |
| **Apicoplast genome** | **(n=44)** | **0.004** | 0.002 | | 0.038 | **0.021** | 0.019 | | 0.098 | **0.002** | 0.000 | | 0.028 | **0.002** | 0.000 | | 0.045 |

**Supplemental Table S12. Summary statistics of nucleotide diversity among cattle-derived *T. parva* strains, and between cattle and the buffalo-derived strain *T. parva* Buffalo LAWR for known *T. parva* antigens.**

KNOWN ANTIGENS ONLY

| | Buffalo-vs-cattle | | | Among Cattle Strains | | |
|---|---|---|---|---|---|---|
| | $\pi_N$ | $\pi_S$ | $\pi_N/\pi_S$ | $\pi_N$ | $\pi_S$ | $\pi_N/\pi_S$ |
| **Mean** | **0.024** | **0.058** | **0.335** | **0.010** | **0.025** | **0.460** |
| Median | 0.009 | 0.040 | 0.262 | 0.001 | 0.012 | 0.240 |
| Max | 0.210 | 0.258 | 0.987 | 0.109 | 0.170 | 2.630 |
| StdDev | 0.044 | 0.055 | 0.313 | 0.024 | 0.037 | 0.600 |

**Supplemental Table S13. Detection of rapidly evolving genes[1].**

| Product name | Locus tag (for individual genes) | Top $\pi_N$ among cattle | Top $\pi_N$ between Muguga and LAWR | Top $\pi_N/\pi_S$ between Muguga and LAWR |
|---|---|---|---|---|
| 5'-3' exoribonuclease 1 | TpMuguga_02g00165 | | X | |
| ABC transporter | TpMuguga_01g00646<br>TpMuguga_02g00016<br>TpMuguga_02g00951<br>TpMuguga_03g00007<br>TpMuguga_03g00864 | X<br>X<br>X<br>X<br>X | | |
| Adaptin N terminal region | TpMuguga_04g00106 | X | | |
| Ankyrin repeat family protein | TpMuguga_03g00538 | X | | |
| AP2 domain protein | TpMuguga_03g00093 | | | X |
| Archease protein family (MTH1598/TM1083) | TpMuguga_03g00010 | | X | X |
| Beta-Casp domain | TpMuguga_03g00560 | X | | |
| Biotin-requiring enzyme family protein | TpMuguga_03g00320 | | X | X |
| Box C/D snoRNA protein 1 | TpMuguga_03g02410 | | | X |
| Chaperone protein DnaJ | TpMuguga_01g02410 | | X | |
| Choline/ethanolamine kinase | TpMuguga_02g00655 | X | | |
| Chromosome segregation protein Spc25 family protein | TpMuguga_03g00851 | | | X |
| Chymosin | TpMuguga_03g02555 | | X | |
| Cyclin-dependent kinase regulatory subunit family protein | TpMuguga_03g00089 | | X | X |
| Cytokine-induced anti-apoptosis inhibitor 1 Fe-S biogenesis | TpMuguga_01g00461 | | X | X |
| DnaJ domain protein | TpMuguga_02g00414 | | X | |
| DSHCT (NUC185) domain protein | TpMuguga_04g00364 | | X | |
| ELM2 domain protein | TpMuguga_01g00312 | | X | |
| EMG1/NEP1 methyltransferase | TpMuguga_03g00619 | X | X | |
| Epsin-2 | TpMuguga_01g00558 | | X | |
| eRF1 methyltransferase catalytic subunit MTQ2 | TpMuguga_04g00566 | X | | |

| | | | | |
|---|---|---|---|---|
| Eukaryotic glutathione synthase ATP binding domain | TpMuguga_01g00265<br>TpMuguga_01g00264 | X<br>X | | |
| Exonuclease 1 | TpMuguga_04g00145 | X | X | |
| Exosome complex component RRP45 | TpMuguga_01g00013 | | | X |
| GDP dissociation inhibitor family protein | TpMuguga_04g00080 | | X | |
| Haemolysin-III related family protein | TpMuguga_04g00201 | | | X |
| Haloacid dehalogenase-like hydrolase<br>**Haloacid dehalogenase-like hydrolase**<br>**Haloacid dehalogenase-like hydrolase** | TpMuguga_01g01075<br>**TpMuguga_01g01078**<br>**TpMuguga_01g01081** | **X** | X | **X** |
| Histone H3-like centromeric protein CSE4 | TpMuguga_02g00044 | | | X |
| Hypothetical protein | **TpMuguga_03g00263** | 31A<br>40AB<br>5AC<br>16ABC | 21B<br>40AB<br>8BC<br>16ABC | 68C<br>5AC<br>8BC<br>16ABC<br>**X** |
| Leucine carboxyl methyltransferase family protein | TpMuguga_02g00808 | X | X | |
| Metallopeptidase family M24 | TpMuguga_03g00462 | | | X |
| Mitochondrial large subunit ribosomal protein (Img2) family protein | TpMuguga_03g02335 | X | X | |
| Myb-like DNA-binding domain protein | TpMuguga_02g00403 | | X | X |
| NADPH:adrenodoxin oxidoreductase mitochondrial | TpMuguga_02g02525 | | X | |
| Nucleoplasmin family protein | TpMuguga_04g00909 | | X | |
| **p104 - 104 kDa microneme/rhoptry antigen** | **TpMuguga_04g00437** | | | **X** |
| **p32 - Merozoite Antigen** | **TpMuguga_01g01056** | **X** | | **X** |
| **PIM** | **TpMuguga_04g00051** | **X** | **X** | **X** |
| Polyubiquitin | TpMuguga_02g00142 | | | X |
| Pre-mRNA splicing factor family protein | TpMuguga_01g01190 | X | X | X |
| Pre-mRNA-splicing factor of RES complex | TpMuguga_02g00881 | | X | |
| Pre-rRNA-processing protein esf-2 | TpMuguga_01g00309 | | X | |
| Protein kinase domain protein | TpMuguga_02g00630 | | | X |

| | | | | |
|---|---|---|---|---|
| Putative integral membrane protein | | 9A<br>7AB<br>1AC<br>4ABC | 8B<br>7AB<br>3BC<br>4ABC | 13C<br>3BC<br>1AC<br>4ABC |
| Rab-GTPase-TBC domain protein | TpMuguga_04g00105 | X | X | X |
| Reactive mitochondrial oxygen species modulator 1 family protein | TpMuguga_04g00826 | X | | |
| Replication factor RFC1 C terminal domain | TpMuguga_03g00565 | X | X | |
| Ribosomal protein L7/L12 C-terminal domain protein | TpMuguga_03g00328 | | | X |
| RNA polymerase II subunit A C-terminal domain phosphatase | TpMuguga_03g00826 | | X | |
| RNAse P Rpr2/Rpp21/SNM1 subunit domain protein | TpMuguga_01g00916 | | | X |
| S-adenosyl-L-methionine-dependent tRNA 4-demethylwyosine synthase | TpMuguga_01g00125 | | X | |
| Sas10/Utp3/C1D family protein | TpMuguga_01g02625 | | | X |
| SEP domain protein | TpMuguga_03g00300 | X | | |
| Sin3 associated polypeptide p18 (SAP18) family protein | TpMuguga_03g00824 | | X | X |
| SVSP family protein | **TpMuguga_01g01225**<br>**TpMuguga_02g00958** | 9A<br>6AB<br>3AC<br>2ABC | 4B<br>6AB<br>10BC<br>2ABC | 16C<br>3AC<br>10BC<br>2ABC<br>**X**<br>**X** |
| **Tash protein PEST motif family protein** | **TpMuguga_04g00164** | X | X | X |
| Telomere recombination | TpMuguga_03g00474 | X | | |
| **Tp1** | **TpMuguga_03g00849** | | | **X** |
| **Tp2** | **TpMuguga_01g00056** | **X** | **X** | **X** |
| **Tp9** | **TpMuguga_02g00895** | **X** | **X** | **X** |

| | | | | |
|---|---|---|---|---|
| TpHN family protein | TpMuguga_01g00609 | X | | |
| | TpMuguga_01g00615 | | | X |
| | TpMuguga_01g00616 | | | X |
| | TpMuguga_01g00605 | | | X |
| | TpMuguga_01g00610 | | | X |
| | TpMuguga_01g00607 | | | X |
| | TpMuguga_01g00619 | | | X |
| | TpMuguga_01g00602 | | | X |
| Tpr family protein | | 3A | | |
| | | | 1B | |
| | | | | 0C |
| | | 28AB | 28AB | |
| | | | 1BC | 1BC |
| | | 1ABC | 1ABC | 1ABC |
| Trafficking protein particle complex subunit 10, TRAPPC10 family protein | TpMuguga_02g00868 | X | X | |
| Transcription factor/nuclear export subunit protein 2 family protein | TpMuguga_02g00884 | | X | X |
| Translation initiation factor 1A / IF-1 family protein | TpMuguga_01g00597 | | | X |
| Translation initiation factor IF-2 | TpMuguga_01g01188 | | X | |
| | TpMuguga_01g02835 | | | X |
| | TpMuguga_04g00278 | | | X |
| | TpMuguga_04g00279 | | X | X |
| | TpMuguga_04g00280 | | | X |
| Type-2 histone deacetylase 2 | TpMuguga_03g02515 | X | X | |
| U3 small nucleolar RNA-associated protein 6 family protein | TpMuguga_03g00023 | X | | |
| Ubiquitin family protein | TpMuguga_01g00315 | X | | |
| Urm1 (Ubiquitin related modifier) family protein | TpMuguga_02g00852 | | X | X |
| Utp11 protein | TpMuguga_01g00183 | | | X |
| WD domain, G-beta repeat family protein | TpMuguga_01g00545 | X | | |
| Ydr279p protein family (RNase H2 complex component) | TpMuguga_03g00595 | | | X |
| Zinc finger A20 and AN1 domain-containing stress-associated protein 9 | TpMuguga_04g00113 | | | X |
| Zinc finger A20 and AN1 domain-containing stress-associated protein 9 | TpMuguga_04g00117 | | | X |
| Zn-finger in Ran binding protein and others family protein | TpMuguga_04g00822 | X | X | |

[1]Genes encoding the proteins with the highest rate of amino acid polymorphism (top 200 $\pi_N$ values) among cattle strains (Muguga, Marikebuni and Uganda) and between cattle (Muguga) and buffalo (LAWR) strains, and the 200 genes with the highest $\pi_N/\pi_S$, ratio

between cattle (Muguga) and buffalo (LAWR) strains. Rows with an "X" indicate an individual gene within one or multiple classes, where "A", "B", and "C" are used when more than eight genes have the same product name. "A" indicates genes falling in the Top $\pi_N$ among cattle class, "B" indicates Top $\pi_N$ between Muguga and LAWR, and "C" indicates Top $\pi_N/\pi_S$ between Muguga and LAWR. If a gene falls in multiple classes, the respective letters are used.
